# Genome-wide protein-DNA interaction site mapping using a double strand DNA-specific cytosine deaminase

**DOI:** 10.1101/2021.08.01.454665

**Authors:** Larry A. Gallagher, Elena Velazquez, S. Brook Peterson, James C. Charity, FoSheng Hsu, Matthew C. Radey, Michael J. Gebhardt, Marcos H. de Moraes, Kelsi M. Penewit, Jennifer Kim, Pia A. Andrade, Thomas LaFramboise, Stephen J. Salipante, Victor de Lorenzo, Paul A. Wiggins, Simon L. Dove, Joseph D. Mougous

## Abstract

DNA–protein interactions (DPIs) are central to such fundamental cellular processes as transcription and chromosome maintenance and organization. The spatiotemporal dynamics of these interactions dictate their functional consequences; therefore, there is great interest in facile methods for defining the sites of DPI within cells. Here, we present a general method for mapping DPI sites *in vivo* using the double stranded DNA-specific cytosine deaminase toxin DddA. Our approach, which we term DddA-sequencing (3D-seq), entails generating a translational fusion of DddA to a DNA binding protein of interest, inactivating uracil DNA glycosylase, modulating DddA activity via its natural inhibitor protein, and DNA sequencing for genome-wide DPI detection. We successfully applied this method to three *Pseudomonas aeruginosa* transcription factors that represent divergent protein families and bind variable numbers of chromosomal locations. 3D-seq offers several advantages over existing technologies including ease of implementation and the possibility to measure DPIs at single-cell resolution.

## Main

Advances in DNA sequencing have promoted rapid expansion in DPI mapping technologies and their applications. Chromatin immunoprecipation sequencing (ChIP-seq) became an early standard for studying both prokaryotic and eukaryotic systems^1^. In this approach, DPIs are identified through chemical crosslinking of DNA–protein complexes, DNA fragmentation, immunoprecipitation of a DNA binding protein (DBP) of interest, crosslink reversal, DNA purification, and DNA sequencing. Sample preparation is technically challenging and requires approximately one week to implement. More recently, Cut&Run and related technologies have gained popularity as alternatives to ChIP-seq^2, 3^. These techniques offer several advantages relative to ChIP-seq including low starting material quantities that permit single cell measurements, the absence of crosslinking and its associated artifacts, and reduced sequencing with improved signal-to-noise^4–6^.

Although powerful, ChIP-seq and Cut&Run-related approaches are fundamentally ex vivo technologies and cannot capture DPIs in living cells. A method that overcomes this limitation is DNA adenine methyltransferase identification (DamID), where the DBP of interest is fused to DAM and DPI site identification occurs through restriction enzyme or antibody mediated methylation site enrichment^7^. However, the utility of this technique is limited by low resolution (1 kb) owing to the frequency of DAM recognition sites (GATC) and by toxicity resulting from widespread adenine methylation. A second approach that facilitates the mapping of DPIs *in vivo* employs mapping the sites of insertion of so-called self-reporting transposons (SRTs). In this technique, a transposase is fused to the DBP of interest, and DPIs are identified by DNA or RNA sequencing to determine sites of transposon insertion^8, 9^. A major limitation to this approach is that transposon insertions occur at low frequency within individual cells (15-100 events per cell), and thus the technology it is not amenable to single cell studies^8^. Additionally, the accumulation of transposon insertions within a population may cause phenotypic consequences through gene disruption.

Nucleic acid-targeting deaminases are a diverse group of proteins that have found a number of biotechnological applications due to their ability to introduce mutations in DNA or RNA. Fusion of the single-stranded DNA (ssDNA) cytosine deaminase APOBEC to catalytically inactive or nickase variants of Cas9 led to the development of the first precision base editor capable of introducing single nucleotide substitutions (C•G-to-T•A) *in vivo*^10^. This breakthrough technology inspired the repurposing of several other ssDNA and RNA-targeting deaminases as base editing tools, including editors that catalyze A•T-to-G•C substitutions in DNA, and RNA transcript editors that induce C to U or A to I modifications^11^. RNA-targeting deaminases have additionally been employed for the identification of RNA–protein complex sites^12^. As the only deaminase known to act preferentially on dsDNA, the bacterial toxin-derived cytosine deaminase, DddA, is unique. We previously capitalized on this feature to develop DddA-derived cytosine base editors (DdCBEs), composed of DddA–TALE fusions that edit the human mitochondrial genome in a programmable fashion^13^. In the current study, we harnessed the dsDNA-targeting capability of DddA in the development of 3D-seq, a new technique for genome-wide DPI mapping.

In DdCBEs, DddA activity is localized to particular sites on DNA by reconstitution of the enzymatic domain of the toxin (amino acids 1264-1427) from split forms fused to sequence-specific targeting proteins^13^. We envisioned an inverse approach whereby fusion of the intact deaminase domain of DddA, referred to herein as DddA, to DBPs with unknown binding sites could be used to define sites of interaction (Fig. 1a). To test the feasibility of this approach, we selected the candidate DNA binding protein GcsR of *P. aeruginosa*. GcsR is a sigma 54-dependent transcription activator of an operon encoding the glycine cleavage system (*gcvH2*, *gcvP2*, and *gcvT2*) and auxiliary glycine and serine metabolic genes (*glyA2* and *sdaA*)^14^. By analogy with closely related sigma 54-dependent regulators, also referred to as bacterial enhancer binding proteins (bEBPs), glycine binding to GcsR is thought to activate transcription of the operon by triggering conformational changes among subunits bound to three 18-bp tandem repeat binding sites in the *gcvH2* promoter region. RNA-seq analyses of *P. aeruginosa* Δ*gcsR* suggest that the *gcvH2* operon may encompass the only genes subject to direct regulation by GcsR^14^.

**Fig. 1:**
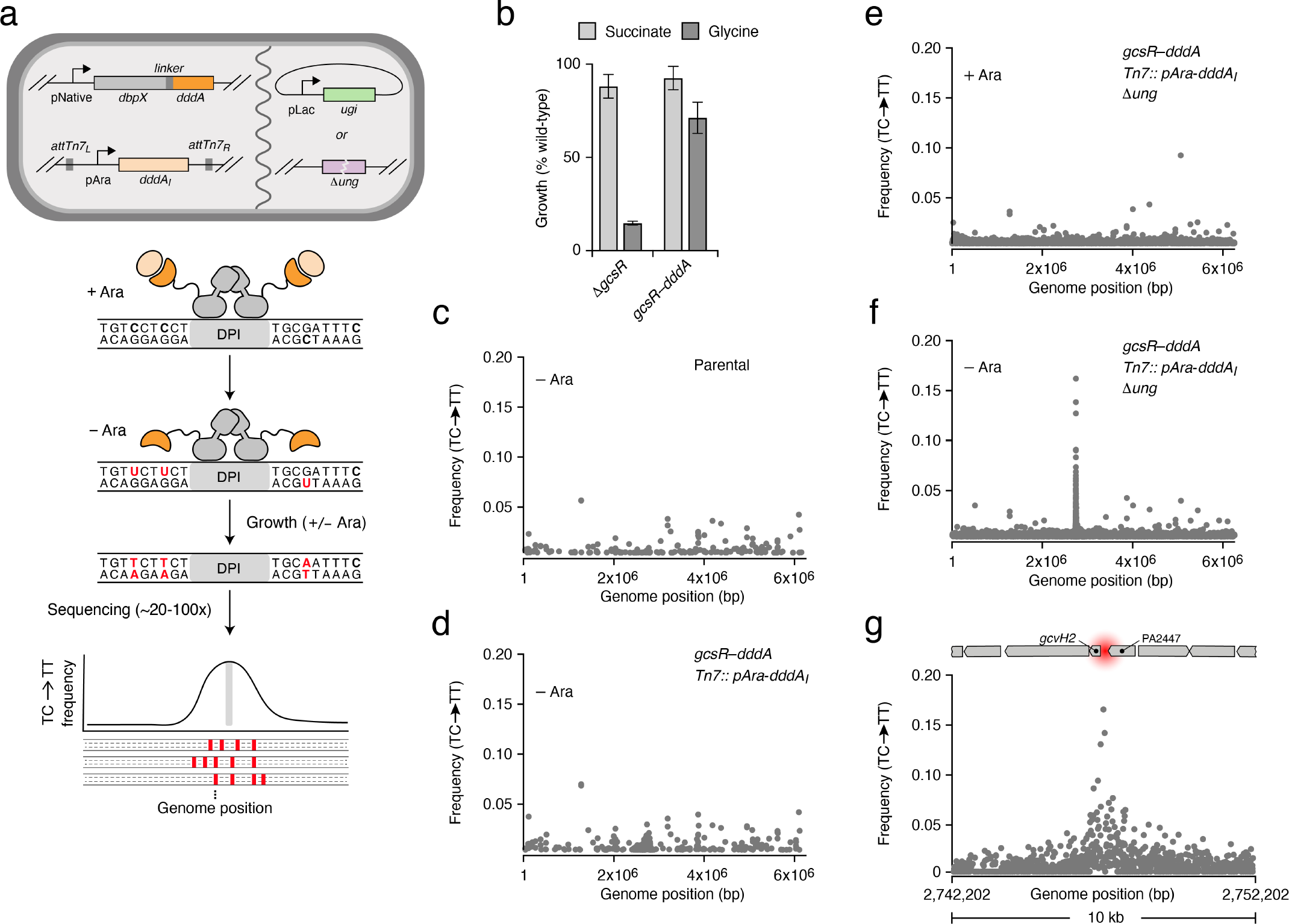
3D-seq is a method for *in vivo* DPI mapping and can be applied to *P. aeruginosa* GcsR. **a,** Diagram providing an overview of the 3D-seq method. Top, cell schematic containing the genetic elements required for 3D-seq. Elements may be integrated into the chromosome or supplied on plasmids. Middle, model depicting localized activity of DddA (dark orange) when fused to a DBP of interest (grey) and after growth in the absence of arabinose to limit production of DddA_I_ (light orange). Bottom, schematized 3D-seq output indicating enrichment of C•G-to-T•A transitions (red) in the vicinity of a DPI site (grey). **b,** Growth yield (normalized to wild-type) of the indicated strains on minimal medium containing glycine or succinate as the sole carbon source. **c-f,** Average (n=4) C•G-to-T•A transition frequency by genome position after passaging cultures of *P. aeruginosa* bearing the indicated genotypes, in the presence or absence of arabinose (Ara) to induce DddA_I_ expression. Data were filtered to remove a prophage hypervariable region and positions with low sequence coverage (<15-fold read depth), and positions with an average transition frequency <0.004 were removed ease of visualization. **g**, Zoomed view of a subset of the data depicted in (**f**). Location of the previously characterized GcsR binding site (red) and adjacent genetic elements shown to scale above.

To capture physiologically relevant DNA binding, we sought to generate a GcsR– DddA translational fusion encoded at the native *gcsR* locus. These efforts revealed that even in the context of fusion to transcription factors under native regulation, DddA exhibits sufficient toxicity to interfere with strain construction. To circumvent this, we inserted the gene encoding the DddA cognate immunity determinant, *dddA_I_*, at the Tn7 attachment site under control of an arabinose inducible promoter (pAra). In this background, and with induction of immunity, we successfully replaced *gcsR* with an open reading frame encoding GcsR bearing an unstructured linker at its C-terminus fused to the deaminase domain of DddA (GcsR–DddA). Activation of the *gcvH2* operon by GcsR is required for *P. aeruginosa* growth using glycine as a sole carbon source^14^. We found that unlike a strain lacking GcsR, strains expressing GcsR–DddA effectively utilize glycine as a growth substrate, suggesting the fusion retains functionality (Fig. 1b).

Our prior work demonstrated that uracil DNA glycosylase (Ung) effectively inhibits uracil accumulation in cells exposed to DddA^15^. Reasoning this DNA repair factor would limit our capacity to detect DddA activity, we deleted *ung* in the GcsR– DddA-expressing strain. Next, we passaged this strain in the presence and absence of arabinose and performed Illumina-based whole genome sequencing (WGS). Data from replicate experiments was minimally filtered to remove positions with low coverage or hypervariability (see methods) and the average frequency of C•G-to-T•A transition events within 5’-TC-3’ contexts were visualized across the *P. aeruginosa* genome (Fig. 1c-f). Other dinucleotide contexts were excluded based on the known strong preference of DddA for thymidine at the -1 position^13^. Remarkably, in samples propagated in the absence of arabinose, we observed a single apparent peak of DddA activity in this minimally filtered data, which was localized to the promoter region of *gcvH2* (Fig. 1f,g). This peak was not observed in samples containing arabinose, nor was it present in parallel studies using a strain containing Ung (Fig. 1d,e).

While a single peak of GcsR::DddA-dependent activity was readily apparent in our minimally processed data, we reasoned that additional filtering to remove background signal would improve the sensitivity and accuracy of our technique. The filters we employed are detailed in the methods and include *i*) accounting for sequencing errors by applying a minimum read count threshold for mutation events (∼1%), *ii*) eliminating positions lacking a neighboring transition event within the approximate length window likely to be accessible to a bound DBP-DddA fusion protein (100 bp), and *iii*) removing transitions representing SNPs present in the parent strain. Most significantly, given our prior observation that modifications catalyzed by free DddA are randomly distributed across genomes, we reasoned that substantial noise reduction could be achieved by removing transitions not reproduced in independent replicates. Visualization of four GcsR–DddA replicate datasets showed that transition events observed in at least three of the samples were highly enriched in the peak region associated with the *gcvH2* promoter (Fig. 2a,b), and therefore this criterion was added to our filtering workflow.

**Fig. 2:**
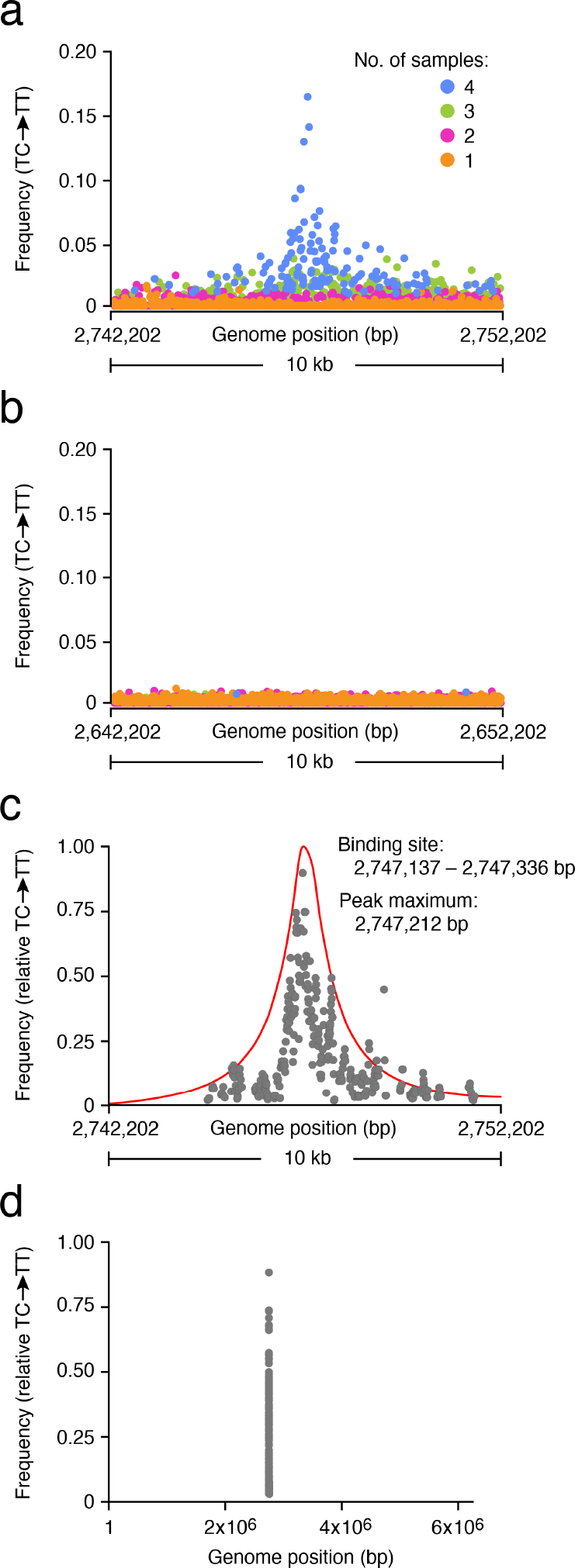
Statistical analyses and data filtering enhance signal-to-noise and allow 3D-seq to precisely map DPIs. **a,b** Average (n=4) C•G-to-T•A transition frequency within the (**a**) primary GcsR 3D-seq peak region or (**b**) a control region located 100,000 bp upstream, with positions colored by the number of replicates in which a transition at that position was observed. **c**, Moving average (75 bp window) of C•G-to-T•A transition frequencies and the curve deriving from our statistical model (red line) calculated from filtered 3D-seq data for the GcsR peak region (see methods). Y-coordinates for the model curve are scaled arbitrarily. **d,** Genome-wide moving average (75 bp window) of C•G-to-T•A transition frequencies calculated for GcsR 3D-seq data after filtering as in (**c**).

In parallel, we sought to develop a statistical analysis able to provide a quantitative means of distinguishing specific DPIs from background noise in 3D-seq data. Our approach employed a null hypothesis test and is described in detail in the methods. Briefly, a null hypothesis consisting of only background enzyme activity was compared to an alternative hypothesis in which a single putative peak was fit by maximum likelihood analysis. The null hypothesis was then either accepted or rejected at a confidence level of 95% using a Generalized Likelihood Ratio Test. If the null hypothesis was rejected, the model containing the peak replaced the null hypothesis and the test was repeated for another putative peak until no more peaks could be detected. P values for each detected peak are estimated and reported (Table S1). The application of these filtering criteria and statistical analyses to our GcsR 3D-seq data dramatically improved the apparent signal-to-noise and placed the major GcsR–DddA binding site centered within the 200 bp region containing the three known binding sites for GcsR^14^ (Fig. 2c,d).

To benchmark the 3D-seq approach, we performed a comparative study using ChIP-seq – a current standard for assessing DPIs genome-wide in bacteria^16^. In place of the *dddA* translational fusion at the 3’ end of *gcsR*, we inserted a sequence encoding the VSV-G epitope to facilitate the necessary immunoprecipitation step of ChIP-seq. Similar to 3D-seq, the most strongly supported candidate binding site for GcsR identified by ChIP-seq localized at the expected region upstream of *gcvH2* (Table S2). We noted in the course of this work that following strain construction, the 3D-seq workflow is considerably streamlined relative to that of ChIP-seq. The hands-on time to process a ChIP-seq sample to the point of sequencing library preparation is approximately one-week in our laboratory, whereas 3D-seq sample preparation constitutes only a genomic DNA preparation that occupies a portion of one day and requires little training.

Given that our initial experiment for detecting GcsR–DddA-catalyzed mutagenesis involved growth for multiple passages, we examined whether a peak of C•G-to-T•A transition frequency in the vicinity of the GcsR binding site could be detected after a shorter period of growth. In continuously growing cultures of *P. aeruginosa* Δ*ung* expressing GcsR–DddA in the absence of DddA_I_ induction we observed a small peak at 9 hrs of propagation and robust DddA–GcsR activity was detected at 20 hrs of growth (Fig. S1). This latter incubation period was thus implemented for subsequent experiments.

We found that Ung inactivation is critical for the detection of GcsR-DNA interactions by 3D-seq (Fig. 1e,f). As an alternative to a *ung* knockout, we considered whether expression of the Ung inhibitor protein, UGI, could achieve sufficient Ung inactivation to reveal GcsR DPIs^17^. This approach is potentially advantageous for 3D-seq in organisms that are difficult to modify genetically. To determine whether expression of UGI could substitute for genetic inactivation of *ung*, we supplied *P. aeruginosa* expressing GcsR–DddA and DddA_I_ with a plasmid possessing Ugi under control of the *lac*UV5 promoter to allow orthogonal modulation of DddA_I_ (arabinose) and Ugi (IPTG). As when Ung was inactivated genetically, we found that inhibition of Ung by UGI expression yielded a high significant peak of C•G-to-T•A transition events centered on the known GcsR binding site upstream of *gcvH2* (Fig. S2, Table S1). This peak was not observed in the empty vector control strain.

To begin to probe the versatility of 3D-seq, we next sought to determine whether it could be successfully applied to the mapping of DPIs for a DBP that is structurally and functionally divergent from GcsR. For this analysis, we selected the regulator GacA, which belongs to a large group of transcription factors known as response regulators. Canonically, phosphorylation of these proteins by cognate histidine kinases enhances their interaction with promoter elements, leading to modulation of transcription ^18^. In the case of GacA, phosphorylation by the sensor kinase GacS promotes binding of GacA to the promoter regions of two small RNA genes, *rsmY* and *rsmZ*^19^. GacS is itself regulated by a second sensor kinase, RetS, which strongly inhibits GacS phosphotransfer to GacA^20^. To further evaluate the capacity of 3D-seq to capture the effects of posttranslational regulation of a transcription factor, we performed our studies in both *ΔgacS* and *ΔretS* backgrounds of *P. aeruginosa*.

During preliminary testing of the 3D-seq protocol with GacA, we found that repressing DddA_I_ production by removing arabinose did not lead to detectable DddA activity. We reasoned that leaky expression of DddA_I_, which is well documented to occur from pAra in *P. aeruginosa*, might be itself sufficient to effectively inhibit DddA in this instance. After exploring alternative promoters without success, we tested a DddA_I_ mutant in which the interaction with DddA is weakened by a C-terminal FLAG epitope fusion (DddA_I–F_, Fig. S3). At high arabinose levels, DddA_I–F_ provided sufficient protection against DddA to permit strain construction and under lower arabinose levels, DddA-dependent C•G-to-T•A transitions were observed.

Consistent with prior studies, 3D-seq revealed GacA binding sites upstream of *rsmY* and *rsmZ* in the Δ*retS* background of *P. aeruginosa* (Fig. 3a-c, Table S1). These peaks were the only significant GacA bindings sites detected and they were not found in the Δ*gacS* strain (Fig. 3d, Table S1). Huang et al. recently reported GacA binds 1125 sites across the *P. aeruginosa* genome, as measured by ChIP-seq^21^. Given the large discrepancy between this result and our findings by 3D-seq, we performed ChIP-seq analysis of GacA in-house. Rather than over-express GacA, which was the strategy adopted by Huang et al., we introduced an epitope-tagged allele of the regulator at its native locus in the Δ*retS* background of *P. aeruginosa*. Consistent with our 3D-seq results and an earlier ChIP-ChIP study^22^, this approach identified regions upstream of *rsmY* and *rsmZ*, enriched 215- and 212-fold, respectively, as the two major bindings sites of GacA (Table S2). A third site located in the promoter region of PA4648 was the only additional site that surpassed our three-fold enrichment significance cut-off. These results added to our confidence in 3D-seq-based DPI site identification and they showed that the methodology can be applied to regulators of different binding modalities and with multiple interaction sites. Finally, they show that 3D-seq can potentially be used to assess PDI dynamics under different regulatory states.

**Fig. 3:**
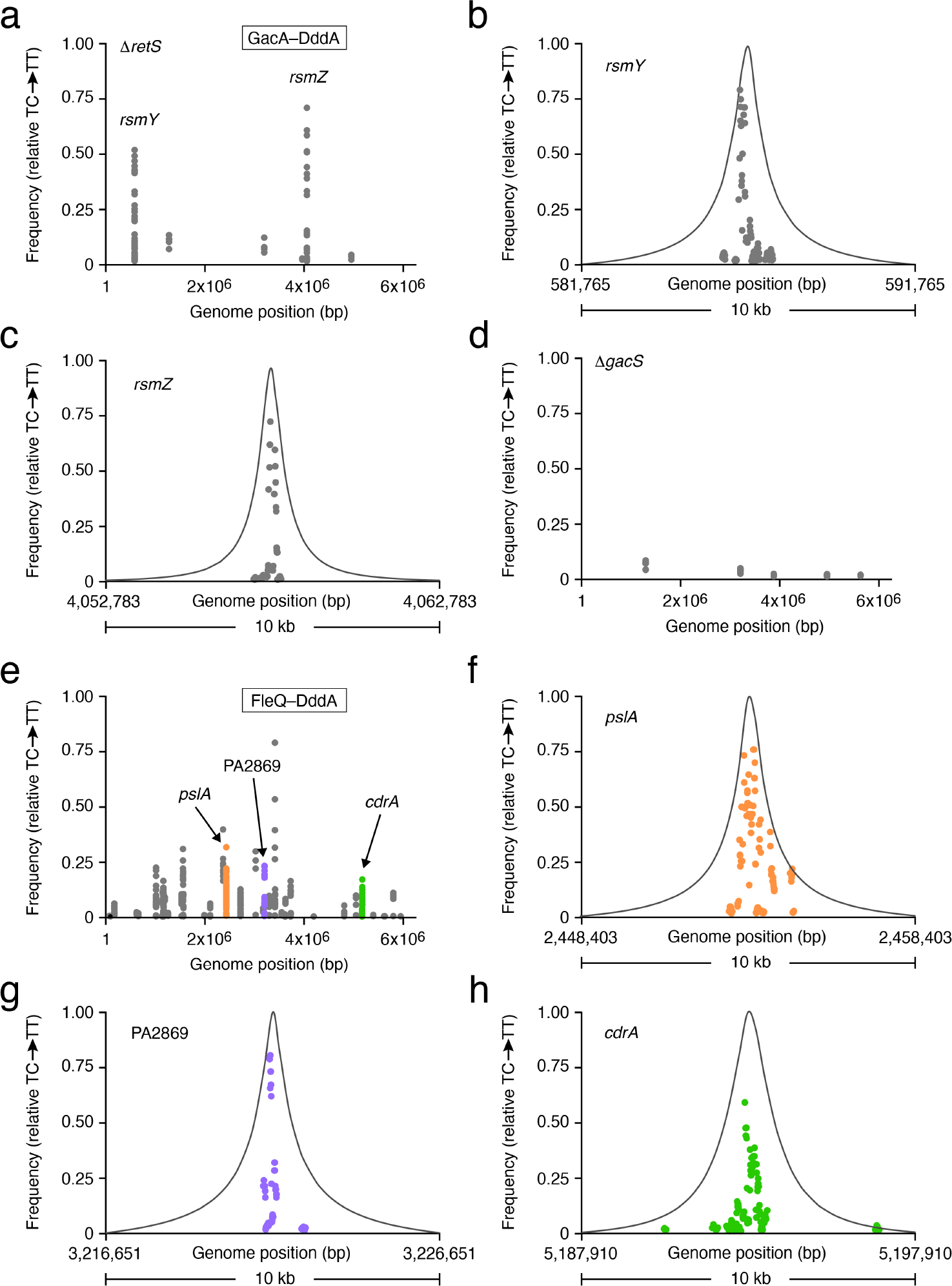
3D-seq maps DPIs for *P. aeruginosa* transcription factors belonging to different families and with varying numbers of binding sites. **a-h,** Moving average (n=4, 75 bp window) of C•G-to-T•A transition frequencies calculated from filtered 3D-seq data deriving from the indicated *P. aeruginosa* strains expressing GacA–DddA (**a-d**) or FleQ–DddA (**e-h**) grown with 0.0005% w/v arabinose for induction of DddA_I–F_. Genome-wide (**a,d,e**) and zoomed (**b,c, f-h**) regions of the data shown in (**a**) or (**e**) are provided. Curves deriving from our statistical model (grey line) calculated from filtered 3D-seq data are shown in the zoomed regions. Y-coordinates for the model curves are scaled arbitrarily. Points in (**f-h)** are colored as in (**e)**.

Although they represent different transcription factor families, our findings show that GcsR and GacA both interact with a limited number of sites on the *P. aeruginosa* chromosome. To gauge the performance of 3D-seq when applied to a DBP with many predicted sites of interaction, we selected the regulator FleQ. This protein is an unusual member of the bEBP family, as it can act as both an activator and repressor, it regulates transcription from both *σ*^54^ and *σ*^70^-dependent promoters, and its regulatory functions appear to be modulated by interaction with an additional protein that does not bind DNA directly, FleN ^23–26^. In its capacity as a *σ*^54^-dependent transcription activator, studies have shown FleQ binds the promoters of several flagellar gene operons; as a *σ*^70^-dependent regulator, it interacts with binding sites adjacent to or overlapping with transcription start sites for several genes involved in exopolysaccharide biosynthesis and can serve as both a repressor and activator depending on availability of the second messenger molecule cyclic-di-GMP^23, 25, 27^. To date, there are no published studies describing the full complement of genes directly regulated by FleQ in *P. aeruginosa.* FleQ was included in the study referenced above that utilized over-expressed transcription factors, but a list of FleQ sites was not provided, and our GacA ChIP-seq and 3D-seq results suggest the general workflow adopted by the authors is problematic^21^.

3D-seq analysis employing FleQ–DddA expressed from its native promoter identified 14 peaks with a significantly elevated frequency of C•G-to-T•A transition events (Fig 3e-h, Table S1). Many of these peaks were localized to previously identified FleQ binding sites. Consistent with expectations for *P. aeruginosa* growing exponentially in liquid media, these included sites upstream of both exopolysaccaride biosynthesis and cell autoaggregation genes known to be repressed by FleQ (e.g. *pelA*, *pslA, siaA*) and flagellar motility genes known to be activated by the protein (e.g. *flhF, fliL, motD*) ^23, 25, 27^. Interestingly, we also identified significant peaks upstream of several uncharacterized genes, including a homolog of the motility gene *fimV* (PA3340), a gene encoded upstream of a c-di-GMP biosynthetic enzyme (PA2869), and a gene with no predicted links to other FleQ-regulated functions (PA3440) (Table S1). These results illustrate the capacity of 3D-seq to sensitively and specifically identify DPIs for proteins that bind at many sites across the genome.

To our knowledge, 3D-seq represents the first method for high-resolution genome-wide recording of DPIs in living cells. In addition to this unique capability of 3D-seq, we found the method offers several advantages over commonly employed technologies for DPI mapping. Key among these is its ease in implementation. Once the appropriate genetic elements are in place, which can in principle be reduced to transformation by a single plasmid, 3D-seq involves simply growing a small volume of the strain under examination followed by genomic DNA preparation and standard WGS. In contrast, ChIP-seq requires a number of specialized reagents, including highly purified antibodies targeting the DBP of interest or an associated epitope tag, and the subsequent technically demanding immunoprecipitation procedure requires several days to complete^1^. Another distinct advantage of 3D-seq is the minimal starting material required. Owing to handling challenges and sample loss occurring at each step of the ChIP-seq protocol, these experiments must generally be initiated with ∼40-80 mL of bacterial culture ^28, 29^. The lower limit on material for a 3D-seq study is defined only by the terminal DNA sequencing technology being utilized. Indeed, we predict that in many circumstances, the genome of a single cell would be adequate for revealing DPIs by 3D-seq.

As performed in this study, 3D-seq exploits the small size of bacterial genomes to cost-effectively obtain high coverage (>100-fold) that can be translated into semi-quantitative measures of DBP occupancy. In eukaryotic organisms with substantially larger genomes, an approach such as this is impractical and enrichment strategies are preferred. Nevertheless, we anticipate that 3D-seq will find application in organisms with large genomes. If experiments are conducted in a manner that permits mutations introduced by the DBP–DddA fusion of interest to approach 100% frequency in the population, far less sequencing depth is required. In another variation, candidate sites could be amplified by PCR and amplicon sequencing would be used to reveal lower frequency modifications.

Despite the strong performance of 3D-seq, there is ample opportunity for optimizing the technology. The straightforward genetic manipulation of *P. aeruginosa* allowed us to generate chromosomally-encoded DBP–DddA fusions and DddA_I_ expression constructs. These sequences, along with that necessary for Ugi expression, could readily be incorporated into a single plasmid, thus eliminating the need for chromosomal manipulations. Future work will compare the performance of 3D-seq by the two approaches.

The resolution of 3D-seq is currently limited by the frequency of cytosines found in the sequence context preferred by DddA, 5’-TC-3’. In *P. aeruginosa*, this dinucleotide motif occurs on average every 12 bp, allowing sufficient resolution to accurately identify DPI sites. Although the average frequency of 5’-TC-3’ is expected to remain relatively consistent across organisms with varying GC content, within particular genomic regions, the frequency of 5’-TC-3’ could diminish substantially and limit resolution. DddA derivatives or novel dsDNA-targeting deaminases with alternative or relaxed sequence specificity hold great promise as a solution to this limitation of 3D-seq^15^.

While we have demonstrated the utility of 3D-seq for the population-level mapping of DPIs involving bacterial transcription factors during in vitro growth, we envision its unique features will catalyze additional applications of the technology going forward. One such feature is the ability to modulate DddA activity through DddA_I_ expression, which enables 3D-seq to capture a snapshot of the protein-DNA landscape during a fixed period of time. This can be particularly advantageous for identifying DPIs during growth under physiological conditions inaccessible to other mapping methods, such as during colonization of a host. The capacity to inducibly inhibit DddA also raises the intriguing possibility of employing 3D-seq to map DPIs within single cells. In this derivative of the technique, we envision a bacterial population would be grown under a condition of interest in the absence of DddA_I_ expression, and subsequently individual clones would be isolated (e.g. as colonies) from media containing the inducer for DddA_I_. Sequencing of these clones, which contain a mutational record of the activity and location of the DBP of interest, will provide heretofore unobtainable genome-wide insights into cell-cell heterogeneity in DPIs. In summary, we anticipate the simplicity of 3D-seq will greatly improve the accessibility of genome-wide DPI mapping studies and its unique attributes will help usher in a new era of DPI measurements in physiological contexts.

## Methods

### Bacterial strains, plasmids, and growth conditions

Detailed lists of all strains and plasmids used in this study can be found in Tables S3 and S4. All *P. aeruginosa* strains were derived from the sequenced strain PAO1^30^ and were grown on Luria-Bertaini (LB) medium at 37°C supplemented as appropriate with 30 µg ml^-1^ gentamicin, 25 µg ml^-1^ irgasan, 5% (*w/v*) sucrose, 1.0 mM IPTG (isopropyl β-D-1-thiogalactopyranoside),and arabinose at varying concentrations. *Escherichia coli* was grown in LB medium supplemented as appropriate with 15 µg ml^-1^ gentamicin, 50 µg ml^-1^ trimethoprim, and 1% rhamnose. *E. coli* strains DH5*α* was used for plasmid maintenance and SM10 (Novagen, Hornsby Westfield, Australia) HB101 (pRK2103) and S17-1 were used for conjugative transfer.

### Plasmid construction

Details of plasmid construction and primer sequences are provided in Tables S5 and S6. Plasmid pEXG2 was used to make the in-frame deletion constructs pEXG2-Δ*gcsR* as well as the VSV-G insertion constructs pEXG2-GcsR-V and pEXG2-GacA*-*V and the DddA fusion constructs pEXG2-*gcsR::dddA,* pEXG2-*gacA–dddA* and pEXG2-*fleQ– dddA*^31^. Plasmid pEXG2-Δ*gcsR*, was constructed by amplification of ∼400 bp regions of genomic DNA flanking *gcsR*, with primers containing restriction sites, followed by digestion and ligation into pEXG2 that had been digested with the appropriate restriction enzymes. C-terminal VSV-G insertion constructs for GcsR–V and GacA–V were made by amplifying ∼400 bp regions flanking each insertion site using primers that contained an in-frame sequence encoding the VSV-G epitope tag. Constructs for generating DddA fusions encoded a protein in which DddA was fused to the C-terminus via a 32aa linker (SGGSSGGSSGSETPGTSESATPESSGGSSGGS). To generate these constructs, primers with 3’ overlapping regions were used to amplify both the linker and *dddA*, as well as 500 bp regions flanking the C-terminus of each gene. Gibson assembly^32^ was then used for the generation of the pEXG2 plasmids containing each construct, and assembly mixes were transformed into *E. coli* DH5*α* expressing DddA_I_ from pSCrhaB2-DddA_I_ to avoid DddA-mediated toxicity. Construction of pEXG2-derived plasmids for deletion of *gacS, retS* and *ung* was previously described^15, 33, 34^. Site-specific chromosomal insertions of the immunity gene *dddA_I_* (with or without a FLAG tag at encoded at the C-terminus) were generated using pUC18T-miniTn7T-Gm-p_BAD_-*araE*. The genes encoding DddA_I_ or DddA_I_-FLAG were amplified and cloned into the KpnI/HindIII sites of this vector through Gibson assembly, to generate pUC18-miniTn7T-Gm-p_BAD_-araE-*dddA_I_* and pUC18T-miniTn7T-Gm-p_BAD_-araE-*dddA_I_-FLAG*.

### Strain construction

*P. aeruginosa* strains containing in-frame deletions of *gcsR, ung, retS or gacS* were constructed by allelic replacement using the appropriate pEXG2-derived deletion construct and were verified by PCR and site specific or genomic sequencing as described previously^31^. *P. aeruginosa* cells synthesizing GcsR with a C-terminal VSV-G epitope tag from the native chromosomal location were made by allelic replacement using vector pEXG2-GcsR-V. *P. aeruginosa* Δ*retS* mutant cells synthesizing GacA with a C-terminal VSV-G epitope tag from the native chromosomal location (*P. aeruginosa* Δ*retS* GacA-V) were made by allelic replacement using vector pEXG2-GacA-V. The *P. aeruginosa* Δ*gcsR*, GcsR-V, and Δ*retS* GacA-V strains were verified by PCR and production of the GcsR-V and GacA-V fusion proteins was verified by Western blotting using an antibody against the VSV-G epitope tag. *P. aeruginosa* strains producing DddA fusion proteins were generated by first engineering the parent strain to express DddA_I_ or DddA_I_-FLAG from the chromosome under arabinose-inducible control by introduction of pUC18T-miniTn7T-Gm-p_BAD_-araE-*DddA_I_* or pUC18T-miniTn7T-Gm-p_BAD_-araE-*DddA_I_-FLAG* and helper plasmids pTNS3 and pRK2013 via tetraparental mating ^35^. After chromosomal integration the GmR marker was removed from these cassettes by Flp/FRT recombination using plasmid pFLP2, which was then cured by sucrose counterselection^36^. *P. aeruginosa* strains synthesizing GcsR-DddA, GacA-DddA or FleQ-DddA from the native chromosomal loci of each regulator were then generated by two-step allelic exchange using the relevant pEXG2 construct. Rhamnose (0.1%, for *E. coli*) or arabinose (0.1%, for *P. aeruginosa*) were maintained during the DddA-fusion expressing strain construction process to minimize DddA toxicity and off-target activity. Fusion-expressing strains were verified by PCR and by assembly of complete genome sequences obtained during 3D-seq analyses.

### Assessing the functionality of the GcsR-DddA fusion protein

To determine the functionality of the GcsR-DddA fusion protein cells were grown in biological triplicate in No Carbon E (NCE) minimal media^37^ containing arabinose (1%) and glycine (20 mM), or arabinose (1%) and succinate (20 mM), at 37°C with aeration for 48 hours. Growth was determined by measuring the culture OD_600_.

### 3D-seq sample preparation and sequencing

#### Culturing of DddA-fusion expressing strains

To generate genomic DNA for 3D-Seq analysis, strains carrying specific DddA fusion constructs and attTn7::araC-P_BAD_-*dddI* (GcsR) or attTn7::araC-P_BAD_-*dddI-FLAG* (GacA, FleQ) were grown for varying amounts of time and with variable levels of arabinose to induce DddA_I_ or DddA_I_-FLAG expression and/or IPTG to induce UGI production from pPSV39-UGI. In each case, the strains were initially streaked for single colonies on LB containing 0.1% or 1% arabinose, and single colonies were used to inoculate quadruplicate liquid cultures containing 0.1% or 1% arabinose. After ∼16 hrs growth, these cultures were then washed with LB and used to inoculate fresh cultures. For GcsR-DddA in *Δung* and *ung*+ backgrounds and for the *Δung* strain without a dddA fusion contruct, washed cultures were inoculated into LB containing 0.1% (negative control) or no (experimental) arabinose at OD_600_ = 0.02, then grown for 8hrs before diluting back to OD_600_ = 0.02. After an additional ∼16 hrs, cultures were again washed and diluted to OD_600_ = 0.02, then grown a final 8 hrs before samples were collected for genomic DNA preparation. For *gacA-dddA* (with Δ*retS* or Δ*gacS*) and *fleQ-dddA*, washed cultures were inoculated into LB containing 0.0005% arabinose at OD_600_ = 0.02, then grown for 6.5 hrs before samples were collected for genomic DNA preparation.

### Genomic DNA preparation and sequencing

Genomic DNA was isolated from bacterial pellets using DNEasy Bood and Tissue Kit (Qiagen). Sequencing libraries for whole-genome sequencing were prepared from 200-300 ng of DNA using DNA Prep Kit (Illumina), with KAPA HiFi Uracil+ Kit (Roche) used in place of Enhanced PCR Mix for the amplification step. Libraries were sequenced in multiplex by paired-end 150-bp reads on NextSeq 550 and iSeq instruments (Illumina).

### ChIP-Seq sample preparation and library construction

200 mL cultures of the *P. aeruginosa* GcsR-V, wild-type, Δ*retS* and Δ*retS* GacA-V strains were grown in biological triplicate to an OD_600_ of 1.5 in LB at 37°C with aeration. 80 mL of culture was crosslinked with formaldehyde (1%) for 30 minutes at room temperature with gentle agitation. Crosslinking was quenched by the addition of glycine (250 mM) and cells were incubated at room temperature for 15 minutes with gentle agitation. Cells were pelleted by centrifugation, washed three times with phosphate buffered saline, and stored at -80°C prior to subsequent processing. Cell pellets were resuspended in 1 mL Buffer 1 (20 mM KHEPES, pH 7.9, 50 mM KCl, 0.5 mM dithiothreitol, 10% glycerol) plus protease inhibitor (complete-mini EDTA-free (Roche); 1 tablet per 10 mL), diluted to a total volume of 5.2 mL and divided equally among four 15 mL conical tubes (Corning). Cells were subsequently lysed and DNA sheared in a Bioruptor water bath sonicator (Diagenode) by exposure to two 8-minute cycles (30 seconds on, 30 seconds off) on the high setting. Cellular debris was removed by centrifugation at 4°C for 20 minutes at 20,000 xg. Cleared lysates were adjusted to match the composition of the immunoprecipitation (IP) buffer (10 mM Tris-HCl, pH 8.0, 150 mM NaCl, 0.1% NP-40 alternative (EMD-Millipore product 492018). The adjusted lysates were combined with anti-VSV-G agarose beads (Sigma) that had been washed once with IP buffer and reconstituted to a 50/50 bead/buffer slurry. For IP, 75 µL of the washed anti-VSV-G beads were added to each of the four aliquots for a given sample. IP was performed overnight at 4°C with gentle agitation. Beads were then washed 5 times with 1 mL IP buffer and 2 times with 1X TE buffer (10 mM Tris-HCl, pH 7.4, 1 mM EDTA). Immune complexes were eluted from beads by adding 150 µL of TES buffer (50 mM Tris-HCl pH 8.0, 10 mM EDTA, 1% Sodium Dodecyl Sulfate (SDS)) and heating samples to 65°C for 15 minutes. Beads were pelleted by centrifugation (5 minutes at 16,000xg) at room temperature and a second elution was performed with 100 µL of 1X TE + 1% SDS. Supernatants from both elution steps were combined and incubated at 65°C overnight to allow cross-link reversal. DNA was then purified with a PCR purification kit (QIAGEN), eluted in 55 µL of 0.1X Elution Buffer and quantified on an Agilent Bioanalyzer. ChIP-Seq libraries were prepared from 1-40 ng of DNA using the NEBNext Ultra II DNA Library Prep Kit for Illumina (NEB). Adaptors were diluted 10-fold prior to ligation. AMPure XP beads (Beckman Coulter) were used to purify libraries, which were subjected to 7 rounds of amplification without size selection. Libraries were sequenced by the Biopolymers Facility (Harvard Medical School) on an Illumina HiSeq2500 producing 75-bp paired-end reads^38^.

### ChIP-Seq data analysis

ChIP-Seq data were analyzed as described previously^38^. Paired-end reads corresponding to fragments of 200 bp or less were mapped to the PAO1 genome (NCBI RefSeq NC_002516) using bowtie2 version 2.3.4.3 ^39^. Only read 1 from each pair of reads was extracted and regions of enrichment were identified using QuEST version 2.4 ^40^. Reads collected from the PAO1 replicates (i.e. IP from PAO1 cells that do not synthesize any VSV-G tagged protein) were merged and served as the mock control for the reads from each of the PAO1 GcsR-V replicates. Merged reads from the PAO1 Δ*retS* replicates served as the mock control for the reads from the PAO1 Δ*retS* GacA-V replicates. The mock control data were used to determine the background for each corresponding ChIP biological replicate. The following criteria were used to identify regions of enrichment (peaks): (i) they must be 3.5-fold enriched in reads compared to the background, (ii) they are are not present in the mock control, (iii) they have a positive peak shift and strand correlation, and (iv) they have a q-value of less than 0.01. Peaks of enrichment for GcsR-V and GacA-V were defined as the maximal region identified in at least two biological replicates. Data were visualized using the Integrative Genomics Viewer (IGV) version 2.5.0^41^. Peak analyses used BEDtools version 2.27.1.

### 3D-seq data analysis

Fastq reads were first pre-processed using the HTStream pipeline v. 1.3.0 (https://s4hts.github.io/HTStream/), where the chain of programs is hts_SuperDeduper -> hts_SeqScreener -> hts_AdapterTrimmer -> hts_QWindowTrim -> hts_LengthFilter -> hts_Stats. In each case logging was enabled and default settings were used, with the following exceptions: 1) For hts_QWindowTrim a window size of 20bp was used with a minimum quality score of 10. 2) For hts_LengthFilter the minimum length was set to half the mean read length. Reads were subsequently aligned to the PAO1-UW reference sequence (https://www.ncbi.nlm.nih.gov/nuccore/NC_002516.2) using Minimap2 v. 2.17-r974-dirty (https://lh3.github.io/minimap2/) and the alignments were saved into sorted BAM files with SAMTools v. 1.10 (https://www.htslib.org/). Alignment position counts were then enumerated using PySAM v. 0.16.0.1 (https://pysam.readthedocs.io/en/latest/) using these settings: read_callback=’alĺ, quality_threshold=20. The reference genome was then surveyed using Biopython v. 1.78 (https://biopython.org/) to determine the proportion of high-quality read-pairs covering each 5’-TC-3’ site (the preferred DddA target sequence context; Mok et al.) on either strand that showed the alternative sequence 5’-TT-3’ (representing cytidine deamination), and corresponding base counts and allele frequencies were tabulated using Pandas v.1.3.0. (https://pandas.pydata.org/).

To generate minimally filtered datasets, sites with sequence coverage of less than 15 read-pairs for that sample were ignored, as were a set of 52 sites within a phage region known to display hypervariability^42^. Average C•G-to-T•A transition frequency was then calculated using remaining positions for each set of quadruplicate samples per condition. To generate more stringently filtered data, sites with >95% C•G-to-T•A transition frequency in all four replicates of a given sample were considered parental SNPs and were ignored. The mean C•G-to-T•A transition frequency was then calculated for each position at which 3 of 4 replicate samples for a given condition exhibited at least 3 sequencing reads containing the mutation. Finally, positions were excluded for which the nearest neighboring position with an average C•G-to-T•A transition frequency >0 was within more than 100 bp. To generate the representations of the data shown in Fig. 1 and 2, this data was further processed by the calculation of a moving average employing a 75 bp window. For statistical analyses, we used data passing these criteria except we required a minimum of only 1 read to contain a given mutation. Additionally, we removed positions from any single sample with

## Statistical analysis

See supplemental methods.

## Data availability

Sequence data associated with this study is available from the Sequence Read Archive at BioProject PRJNA748760.

## Code availability

Computer code generated for this study is available from GitHub at https://github.com/marade/3DSeqTools.

## Acknowledgements

The authors wish to thank members of the Mougous laboratory for helpful suggestions. This work was supported by the NIH (AI080609 to JDM, AI143771 to S.L.D, GM128191 to P.A.W and DK089507 to SJS), and the Cystic Fibrosis Foundation (SINGH19R0 to SJS and a postdoctoral fellowship to M.J.G.), and the SynBio4Flav (H2020-NMBP-TR-IND/H2020-NMBP-BIO-2018-814650) and MIX-UP (MIX-UP H2020-BIO-CN-2019-870294) contracts of the European Union (to V.L.). E.V. was the recipient of a Fellowship from the Education Ministry, Madrid, Spanish Government (FPU15/04315). JDM is an HHMI Investigator.

## Author contributions

L.A.G, E.V., S.B.P, M.H.D., V.L., S.L.D. and J.D.M designed the study, L.A.G, E.V., J.C.C, F.H., K.M.P., J.K., and P.A.A. performed experiments, L.A.G, E.V., S.B.P, M.C.R., M.J.G., T.L., S.J.S, P.A.W., S.L.D. and J.D.M analyzed data, and S.B.P and J.D.M wrote the manuscript with input from the other authors.

## Author Information

Correspondence and requests for materials should be addressed to J.D.M. or S.L.D.

---

**Figure S1.**
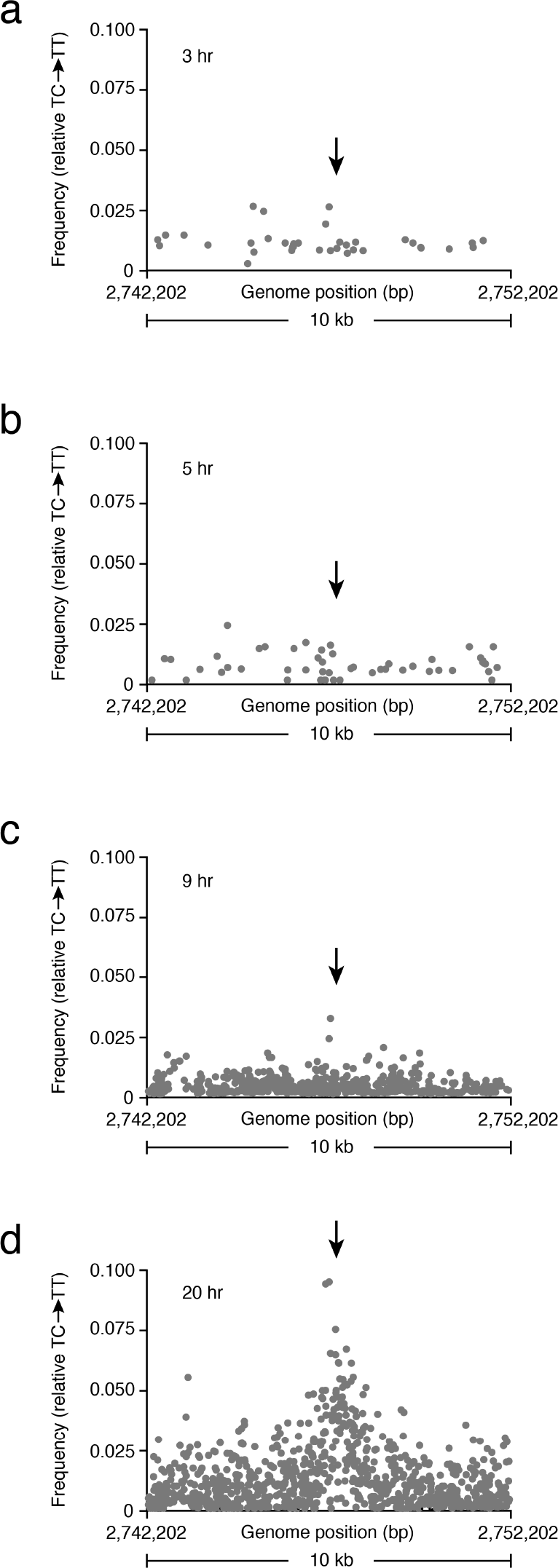
Transition mutations associated with GcsR:DddA activity accumulate over time. **a-d,** Average (n=4) C•G-to-T•A transition frequency within the primary GcsR 3D-seq peak region after the indicated growth period and in the absence of arabinose. Data were filtered as in Fig. 1. The arrow indicates the approximate position of the known GcsR binding site.

**Figure S2.**
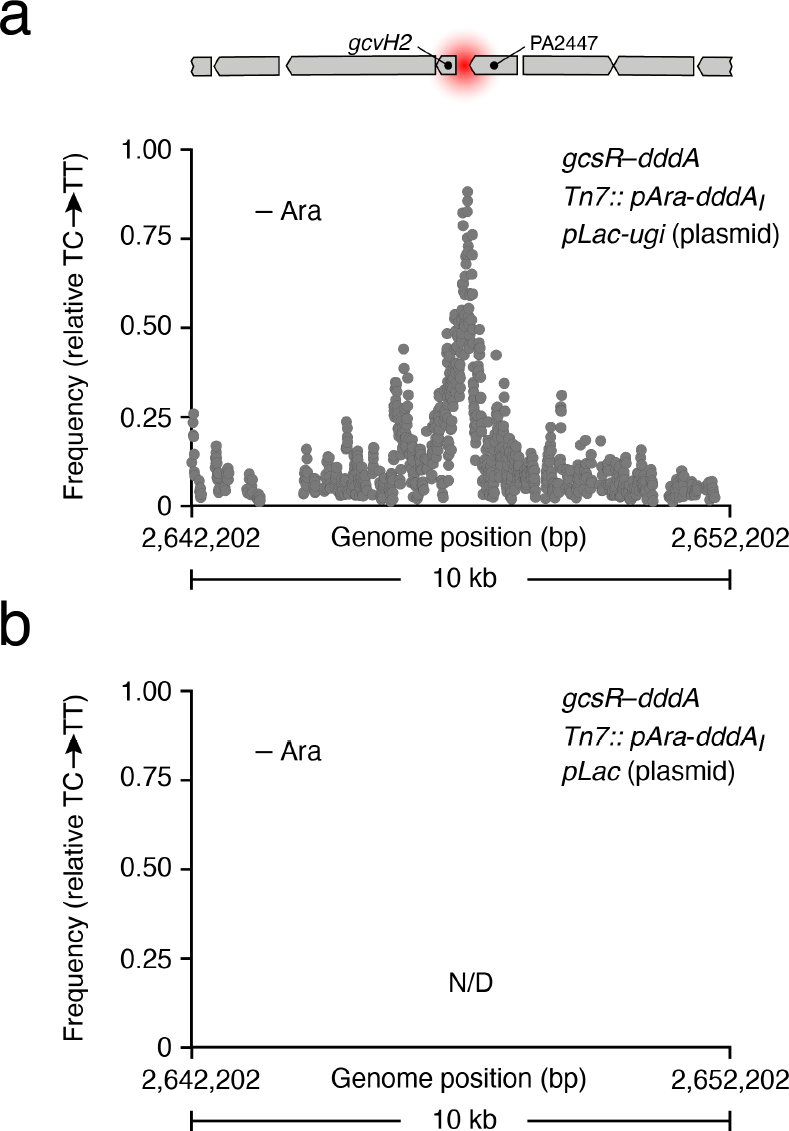
Ugi expression can substitute for genetic inactivation of *ung* in 3D-seq. **a,b,** Moving average (n=4, 75 bp window) of C•G-to-T•A transition frequencies calculated from filtered 3D-seq data deriving from the indicated *P. aeruginosa* strains grown in the absence of arabinose for 20 hrs. IPTG was included to induce the expression of Ugi throughout the growth period. The location of the previously characterized GcsR binding site (red) and adjacent genetic elements are shown to scale above.

**Figure S3.**
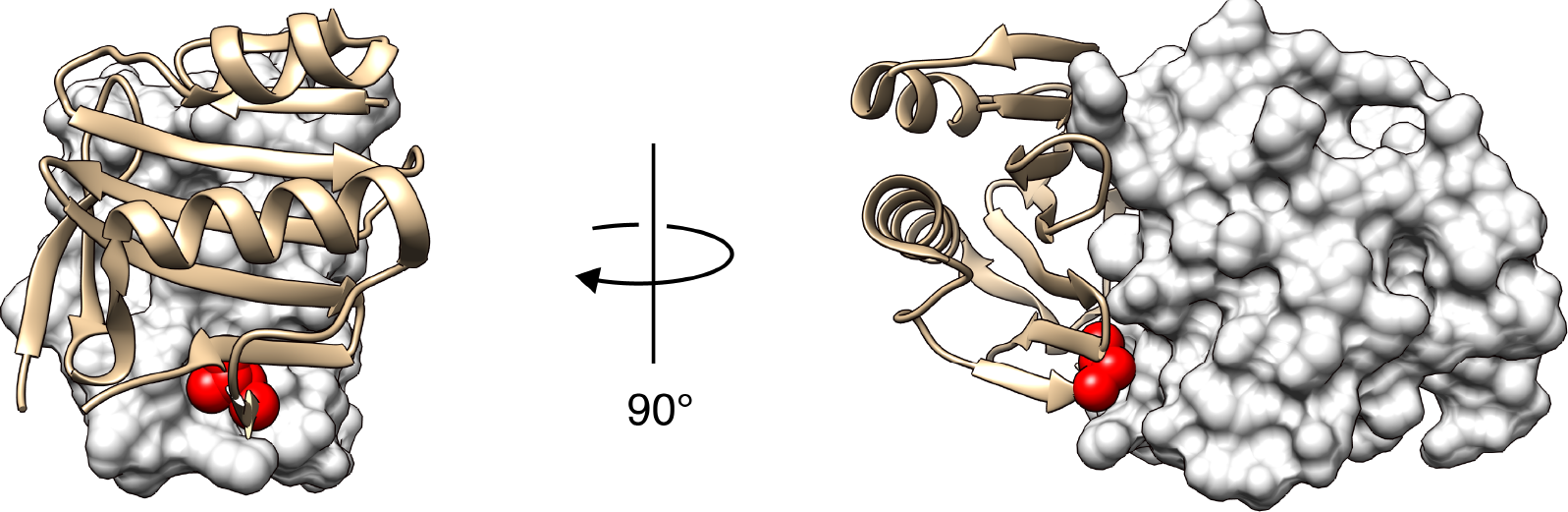
The C-terminus of DddA_I_ abuts DddA. X-ray crystal structure of the DddA_I_–DddA complex in ribbon and surface representation, respectively. The C-terminal amino acid of DddA_I_ (Leu123) is colored red and shown in space filling representation to highlight its position against the surface of DddA.

**Table S1.**
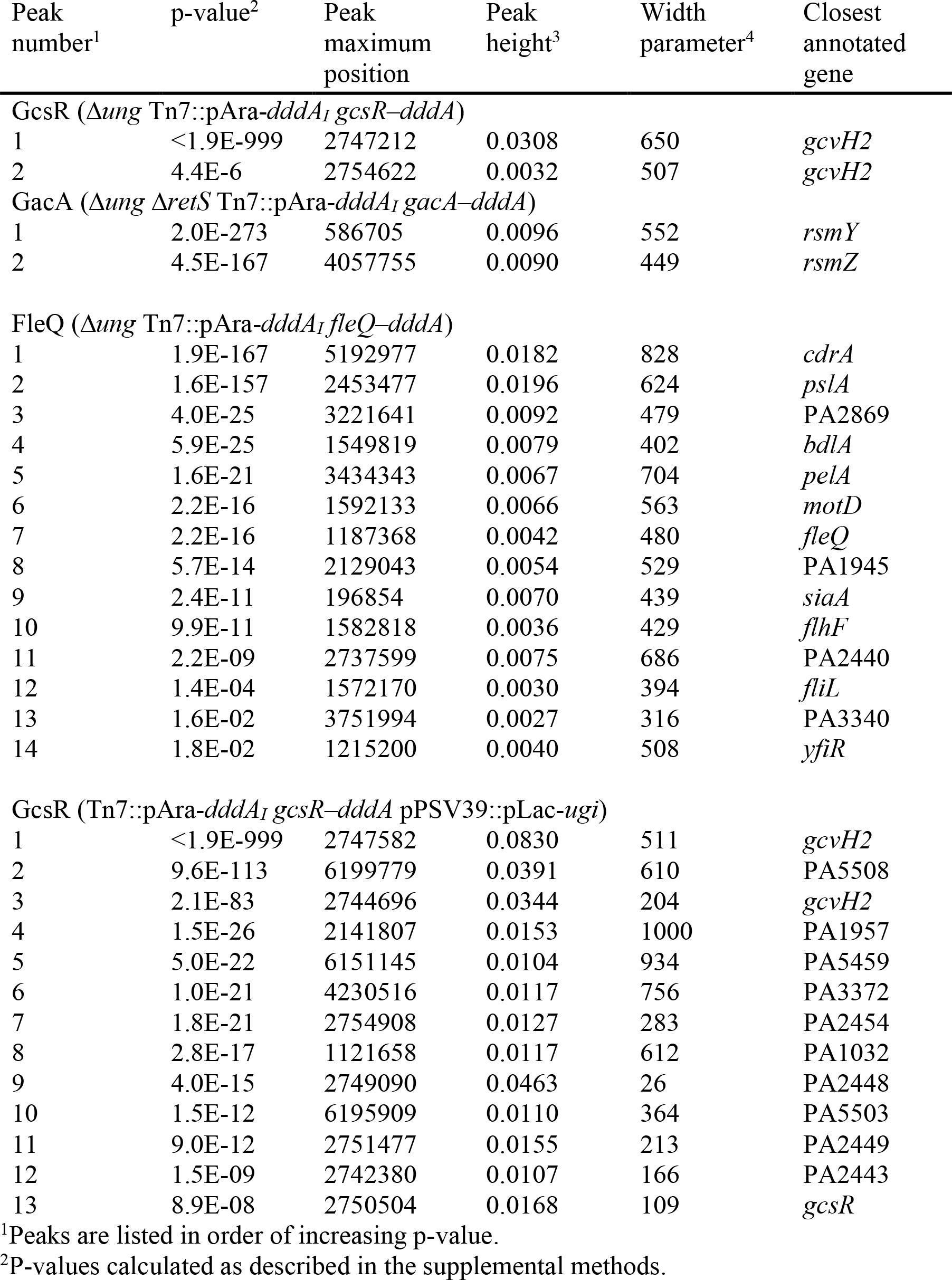
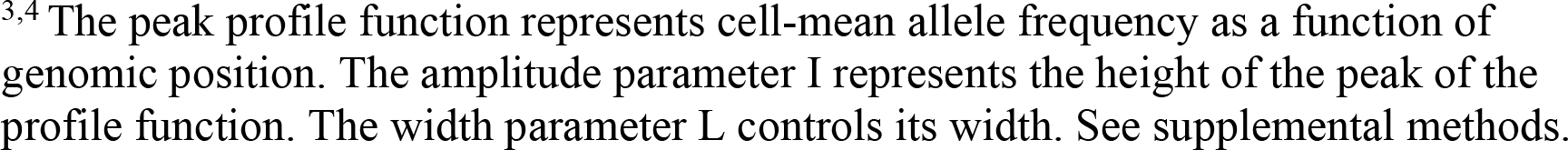
Significant peaks detected in this study by 3D-seq.

| Peak number<sup>1</sup> | p-value<sup>2</sup> | Peak maximum position | Peak height<sup>3</sup> | Width parameter<sup>4</sup> | Closest annotated gene |
| --- | --- | --- | --- | --- | --- |
| **GcsR ($\Delta ung$ Tn7::pAra-*dddA*<sub>I</sub> *gcsR*-*dddA*)** |  |  |  |  |  |
| 1 | <1.9E-999 | 2747212 | 0.0308 | 650 | *gcvH2* |
| 2 | 4.4E-6 | 2754622 | 0.0032 | 507 | *gcvH2* |
| **GacA ($\Delta ung \Delta retS$ Tn7::pAra-*dddA*<sub>I</sub> *gacA*-*dddA*)** |  |  |  |  |  |
| 1 | 2.0E-273 | 586705 | 0.0096 | 552 | *rsmY* |
| 2 | 4.5E-167 | 4057755 | 0.0090 | 449 | *rsmZ* |
| **FleQ ($\Delta ung$ Tn7::pAra-*dddA*<sub>I</sub> *fleQ*-*dddA*)** |  |  |  |  |  |
| 1 | 1.9E-167 | 5192977 | 0.0182 | 828 | *cdrA* |
| 2 | 1.6E-157 | 2453477 | 0.0196 | 624 | *pslA* |
| 3 | 4.0E-25 | 3221641 | 0.0092 | 479 | PA2869 |
| 4 | 5.9E-25 | 1549819 | 0.0079 | 402 | *bdlA* |
| 5 | 1.6E-21 | 3434343 | 0.0067 | 704 | *pelA* |
| 6 | 2.2E-16 | 1592133 | 0.0066 | 563 | *motD* |
| 7 | 2.2E-16 | 1187368 | 0.0042 | 480 | *fleQ* |
| 8 | 5.7E-14 | 2129043 | 0.0054 | 529 | PA1945 |
| 9 | 2.4E-11 | 196854 | 0.0070 | 439 | *siaA* |
| 10 | 9.9E-11 | 1582818 | 0.0036 | 429 | *flhF* |
| 11 | 2.2E-09 | 2737599 | 0.0075 | 686 | PA2440 |
| 12 | 1.4E-04 | 1572170 | 0.0030 | 394 | *fliL* |
| 13 | 1.6E-02 | 3751994 | 0.0027 | 316 | PA3340 |
| 14 | 1.8E-02 | 1215200 | 0.0040 | 508 | *yfiR* |
| **GcsR (Tn7::pAra-*dddA*<sub>I</sub> *gcsR*-*dddA* pPSV39::pLac-*ugi*)** |  |  |  |  |  |
| 1 | <1.9E-999 | 2747582 | 0.0830 | 511 | *gcvH2* |
| 2 | 9.6E-113 | 6199779 | 0.0391 | 610 | PA5508 |
| 3 | 2.1E-83 | 2744696 | 0.0344 | 204 | *gcvH2* |
| 4 | 1.5E-26 | 2141807 | 0.0153 | 1000 | PA1957 |
| 5 | 5.0E-22 | 6151145 | 0.0104 | 934 | PA5459 |
| 6 | 1.0E-21 | 4230516 | 0.0117 | 756 | PA3372 |
| 7 | 1.8E-21 | 2754908 | 0.0127 | 283 | PA2454 |
| 8 | 2.8E-17 | 1121658 | 0.0117 | 612 | PA1032 |
| 9 | 4.0E-15 | 2749090 | 0.0463 | 26 | PA2448 |
| 10 | 1.5E-12 | 6195909 | 0.0110 | 364 | PA5503 |
| 11 | 9.0E-12 | 2751477 | 0.0155 | 213 | PA2449 |
| 12 | 1.5E-09 | 2742380 | 0.0107 | 166 | PA2443 |
| 13 | 8.9E-08 | 2750504 | 0.0168 | 109 | *gcsR* |

<sup>1</sup>Peaks are listed in order of increasing p-value.<sup>2</sup>P-values calculated as described in the supplemental methods.
<sup>3,4</sup> The peak profile function represents cell-mean allele frequency as a function of genomic position. The amplitude parameter $I$ represents the height of the peak of the profile function. The width parameter $L$ controls its width. See supplemental methods.

**Table S2.** Significant peaks detected in this study by ChIP-seq.

| Peak number<sup>1</sup> | Fold enrichment | Peak maximum | Peak start | Peak end | Closest annotated gene |
| --- | --- | --- | --- | --- | --- |
| GcsR (*gcsR*–VSV-G) |  |  |  |  |  |
| 1 | 92 | 2747441 | 2746047 | 2748292 | *gcvH2* |
| 2 | 13 | 5402788 | 5402462 | 5403174 | *lipC* |
| 3 | 9.1 | 945626 | 945487 | 945780 | PA0864 |
| 4 | 5.6 | 2516215 | 2516073 | 2516373 | PA2288 |
| 5 | 3.4 | 4373881 | 4373741 | 4374013 | PA3904 |
| 6 | 3.4 | 3070313 | 3070176 | 3070417 | PA2715 |
| 7 | 3.2 | 1205058 | 1204920 | 1205189 | PA1112a |
| GacA ($\Delta retS$ *gacA*–VSV-G) | | | | | |
| 1 | 22E01 | 586983 | 586145 | 587484 | *rsmY* |
| 2 | 21E01 | 4057808 | 4057399 | 4058161 | *rsmZ* |
| 3 | 3.6 | 5214833 | 5214674 | 5215156 | PA4648 |

<sup>1</sup>Peaks are listed in order of decreasing fold enrichment.

**Table S3:** Bacterial strains used.

| Strain | Source |
| --- | --- |
| ***P. aeruginosa* strains** |  |
| PAO1 | 30 |
| $\Delta ung$ | 15 |
| attTn7::*araC*-P<sub>BAD</sub>-*dddA*<sub>I</sub> | This study |
| $\Delta ung$ attTn7::*araC*-P<sub>BAD</sub>-*dddA*<sub>I</sub> | This study |
| attTn7::*araC*-P<sub>BAD</sub>-*dddA*<sub>I</sub> *gcsR*-*dddA* | This study |
| $\Delta ung$ attTn7::*araC*-P<sub>BAD</sub>-*dddA*<sub>I</sub> *gcsR*-*dddA* | This study |
| attTn7::*araC*-P<sub>BAD</sub>-*dddA*<sub>I</sub> *gcsR*-*dddA* pPSV39(empty) | This study |
| attTn7::*araC*-P<sub>BAD</sub>-*dddA*<sub>I</sub> *gcsR*-*dddA* pPSV39-*ugi* | This study |
| attTn7::*araC*-P<sub>BAD</sub>-*dddA*<sub>I</sub>-FLAG | This study |
| $\Delta ung$ attTn7::*araC*-P<sub>BAD</sub>-*dddA*<sub>I</sub>-FLAG | This study |
| $\Delta ung$ attTn7::*araC*-P<sub>BAD</sub>-*dddA*<sub>I</sub>-FLAG *gacA*-*dddA* | This study |
| $\Delta ung$ attTn7::*araC*-P<sub>BAD</sub>-*dddA*<sub>I</sub>-FLAG *gacA*-*dddA* $\Delta retS$ | This study |
| $\Delta ung$ attTn7::*araC*-P<sub>BAD</sub>-*dddA*<sub>I</sub>-FLAG *gacA*-*dddA* $\Delta gacS$ | This study |
| $\Delta ung$ attTn7::*araC*-P<sub>BAD</sub>-*dddA*<sub>I</sub>-FLAG *fleQ*-*dddA* | This study |
| $\Delta gcsR$ | This study |
| GcsR-V | This study |
| $\Delta retS$ | This study |
| $\Delta retS$ GacA-V | This study |
| ***E. coli* strains** |  |
| DH5α | |
| SM10 | Novagen |
| HB101 | 43 |
| S17-1 | 44 |

**Table S4:** Plasmids used.

| Plasmid | Reference |
| --- | --- |
| pEXG2 | 31 |
| pSCrhaB2 | 45 |
| pPSV39 | 46 |
| pUC18T-miniTn7T-Gm-pBAD-araE | 35 |
| pRK2103 | 47 |
| pTNS3 | 48 |
| pFLP2 | 49 |
| pEXG2-$\Delta ung$ | 15 |
| pEXG2-$\Delta retS$ (pEXG2_ΔPA4856) | 34 |
| pEXG2-$\Delta gacS$ (pEXG2_ΔPA0928) | 33 |
| pSCrhaB2-*dddA<sub>I</sub>* | This study |
| pEXG2-*gcsR-dddA* | This study |
| pEXG2-*gacA-dddA* | This study |
| pEXG2-*fleQ-dddA* | This study |
| pEXG2-$\Delta gcsR$ | This study |
| pEXG2-GscR-V | This study |
| pEXG2-GacA-V | This study |
| pPSV39-*ugi* | This study |
| pUC18T-miniTn7T-Gm-pBAD-araE-*dddA<sub>I</sub>* | This study |
| pUC18T-miniTn7T-Gm-pBAD-araE-*dddA<sub>I</sub>*-FLAG | This study |

**Table S5:**
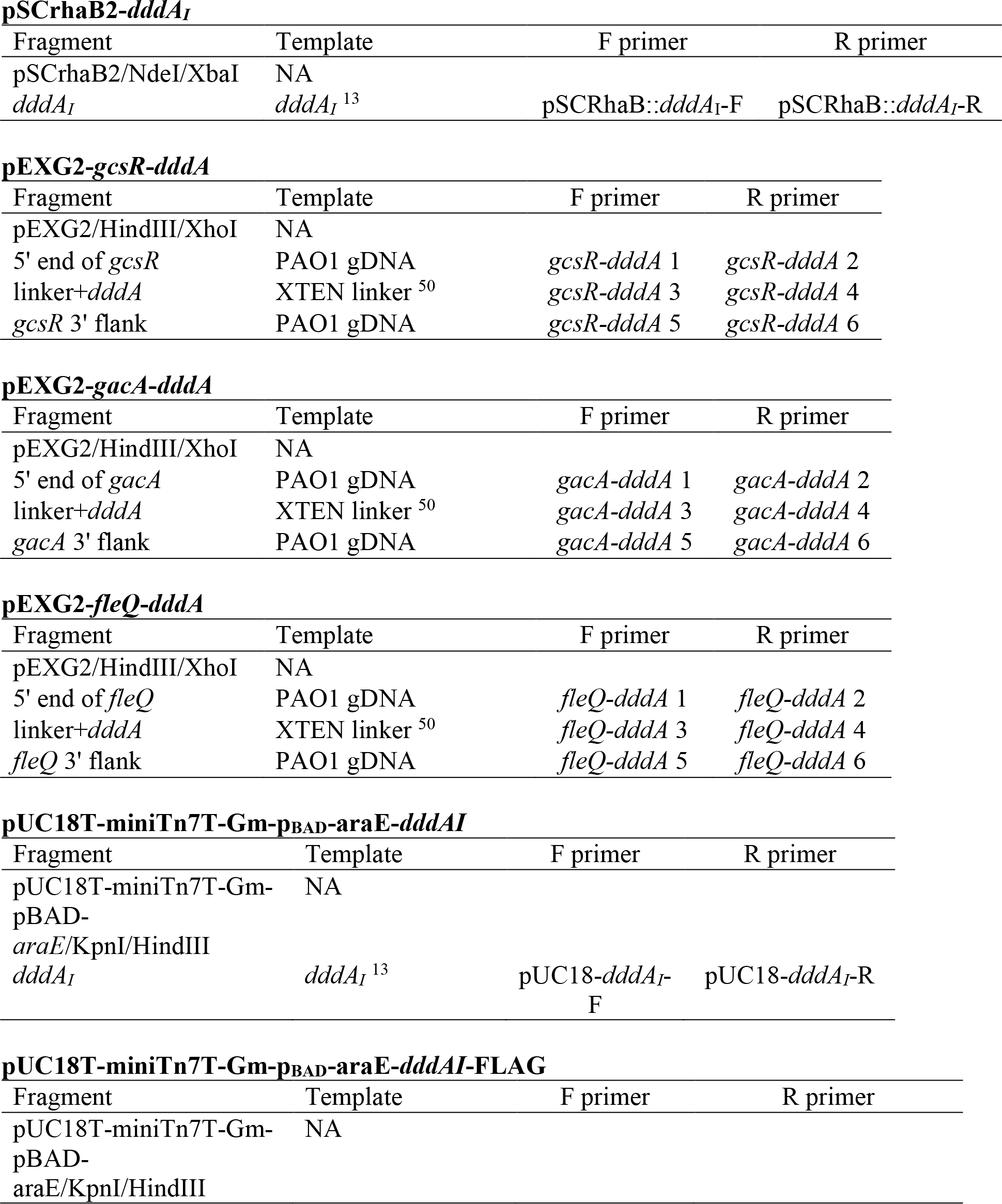
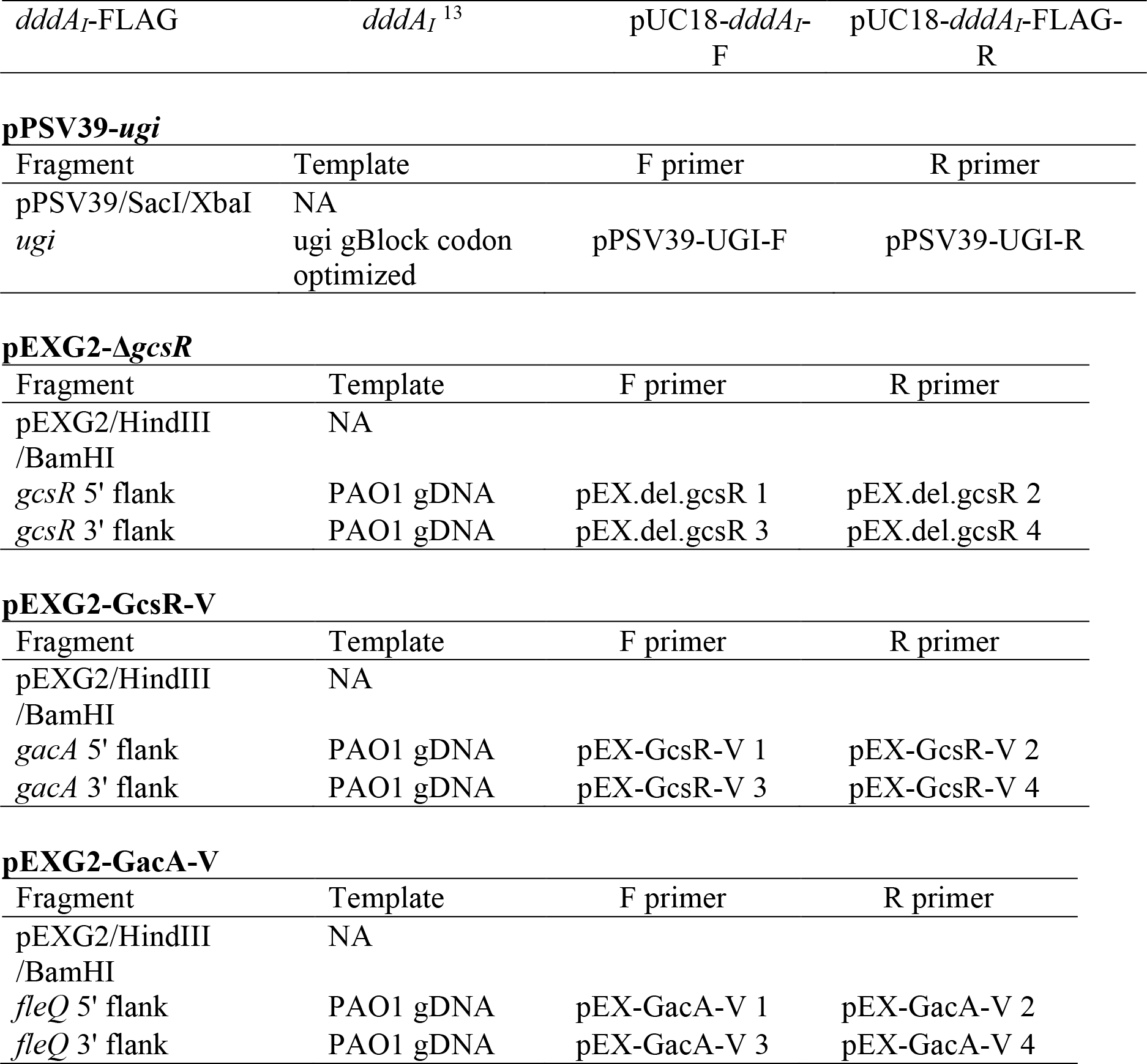
Plasmid construction. The following plasmids were generated by combining the individual fragments listed using either Gibson cloning or overlap extension PCR. Primer sequences are listed in Table S6.

**Table S6:**
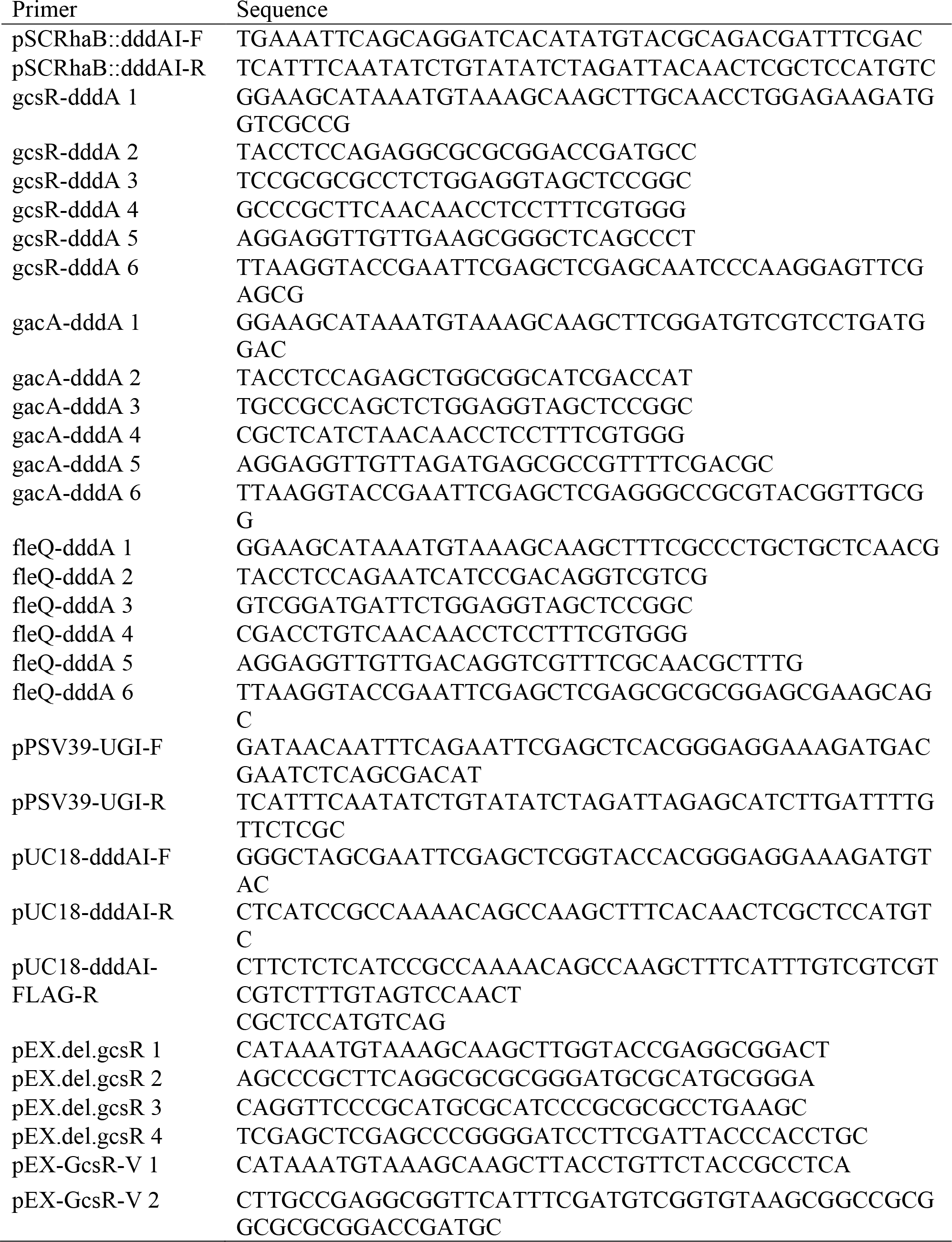
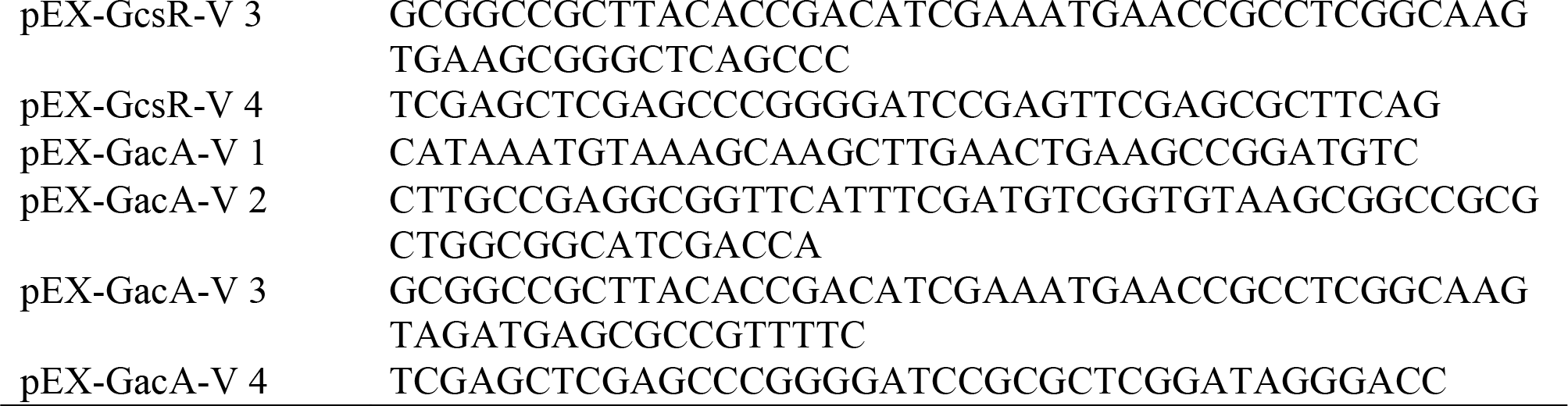
Primer sequences.

