## Supplemental Methods for "Genome-wide protein-DNA interaction site mapping using a double strand DNA-specific cytosine deaminase"

### Supplemental methods
### (Statistical analyses of 3D-seq data)

#### I. STATISTICAL ANALYSIS

We divide the analysis into two steps: peak detection and peak-parameter inference. In the first peak-detection step, we used a canonical frequentist approach: null hypothesis testing [1] to determine the number and approximate position of the peaks in the data. Then, in a second step, we optimized the model parameters describing each peak individually using a slower but more accurate numerical Maximum Likelihood Estimation (MLE) to optimize peak parameter inference.

#### II. BIOPHYSICAL MODEL FOR THE ALLELE FREQUENCY

Motivated by the DNA effective-concentration model (*e.g.* [2]), we modeled the cell-mean allele frequency at locus $j$ as:

$$\overline{r}_j = \mu_0 + \sum_J \delta\mu_J(x_j), \tag{1}$$

where the first term represents the activity on non-localized DddA-transcription-factor fusions and the second term represents the activity, at genomic position $x_j$, for a fusions specifically bound at binding site $J$ at genomic position $\ell_J$. $\delta\mu_J$ will form an allele-frequency peak around site $J$: it will be large when sequence $j$ and site $J$ are proximal and nearly zero for sequences distal to the sites.

For the functional form of the peak profile, we will again consider the DNA effective-concentration model (*e.g.* [2]). We will model the peak profile as a generalized Cauchy function:

$$C(x/L; a, D) \equiv \left[1 + |x/L|^D\right]^{-a/D}, \tag{2}$$

for $D = 1$. In this model $x \equiv x_j - \ell_J$ is the genomic displacement (in bp) between sequence $j$ and binding site $J$. The scaling exponent $a$ is a model parameter that controls the rapidity with which the tails decay away from the peak. Its value is determined by chromatin structure and we expect $1 < a < 1.5$ [2–4]. The parameter $L$ defines the width of the peak and depends both on the structure of the protein fusion as well as chromatin structure. The peak profile is shown in Fig. 1.

In practice, it will be convenient to only consider peak profile functions with local position support. We will therefore use a generalized Cauchy that is cut off at $|x/L| > 16$:

$$C'(\tfrac{x}{L}; a, D) \equiv \begin{cases} \frac{C(\frac{x}{L};a,D) - C(16;a,D)}{C(0;a,D) - C(16;a,D)}, & |x/L| \le 16 \\ 0, & |x/L| > 16 \end{cases}, \tag{3}$$

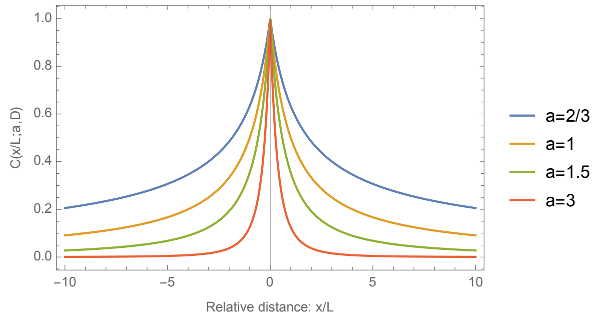


FIG. 1. **The peak profile is modeled by a generalized Cauchy function.** Parameter $a$ controls the power-law decay of the tails with relative genomic distance and parameter $D$ control the geometry of the summit (not shown).

which makes no qualitative difference to shape of the peak profile.

Our model for the mean allele frequency at locus $j$ due to specific binding at site $J$ is therefore:

$$\delta\mu_J(x_j) \equiv \delta\mu(x_j; \theta_J) \equiv I_J C'(\tfrac{x_j - \ell_J}{L_J}; a_J, 1), \tag{4}$$

where the parameter vector contains the following parameters:

$$\theta_J = (I_J, \ell_J, a_J, L_J), \tag{5}$$

and the last undefined parameter $I_J$ controls the peak profile amplitude (or height). The model results in an excellent fit to the observed allele frequency peaks, as shown in Fig. 2.

##### 1. *Modeling of the distribution of allele frequency*

Our model so far describes the cell-mean allele frequency at a genetic locus, however the creation of alleles is a stochastic process. Furthermore, the cells are mutated while the culture is growing, therefore alleles that are created early, grow with the population and have a higher frequency than alleles created late in the growth process. This is the well-known *jackpot phenomenon* [5]. Although this type of principled analysis is possible it would require a great number of parameters and potentially experimental calibrations [6]. Instead, we will implement a much more tractable and practical approach to the modeling of the expected distribution of allele frequencies at a given locus: We will model $r_i$ as a Gaussian random variable with a locus-specific mean (Eq. 1) and variance that is proportional to the mean.

Let the data $D$ be defined as the set of $N$ allele frequencies $r_i$ and genomic positions $x_i$ pairs:

$$D \equiv \{(x_i, r_i)\}_{i=1\ldots N}. \tag{6}$$

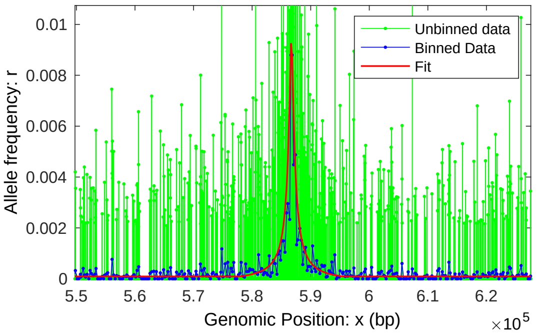


FIG. 2. **A typical fit to a peak.** The unbinned and binned data are compared to the model for the allele frequency for GacA. Although the raw allele frequency appears qualitatively higher than the binned and model frequencies, all three are in fact mutually consistent. The apparent high allele frequencies seen in the raw data are an artifact of the noise in a signal consisting of fluctuations between high allele-frequency values (rare) and zeros (frequent). This noise at the basepair resolution is a consequence of low DddA expression and the finite population size. In contrast both the binned and model allele frequencies can be understood as averaged to reduce this noise: The binned frequency is averaged over a narrow window of genetic loci whereas the model represents the estimated cell-mean allele frequency (i.e. averaged over an infinite-sized population).

We modeled the allele frequencies $r_i$ as a Gaussian random variable. We assume a *locus-dependent* mean $\mu_i$ and variance $\sigma_i^2$:

$$R_i \sim \mathcal{N}(\mu_i, \sigma_i), \tag{7}$$

where $R_i$ is capitalized because it is being interpreted as a random variable, $\mathcal{N}$ is the normal distribution.

#### III. PEAK DETECTION BY NULL HYPOTHESIS TESTING

##### 1. *Data binning and processing for peak detection*

Since the peak features are wide compared to single basepair resolution, it is convenient to detect the peaks initially at low resolution before optimizing the peak parameters using the full resolution data. The central limit theorem guarantees that for sufficiently large bins, the $r_{j'}$ will be normally distributed, simplifying the analysis. However, as the size of the bins grows, so does the noise from the background enzyme activity. We compromised with a bin size of 250 bp.

One important feature of the allele frequency data is that not every base is a target due to the TC-sequence specificity of DddA. We therefore binned the data using a protocol that avoided the introduction of bins weighted by the number of target sites. We divided the genome into 250 bp bins. In each bin $j'$, we have data indexes $j \in \mathcal{J}_{j'}$. We defined the position of the binned target $x_{j'}$ as the weighted average of the sites in that bin:

$$x_{j'} \equiv \frac{\sum_{j\in\mathcal{J}_J} x_j r_j}{\sum_{j\in\mathcal{J}_J} r_j}. \tag{8}$$

If all $r_j$ were zero in the bin, the index position $x_{j'}$ was assigned the mean of $x_j$ for $j \in \mathcal{J}_{j'}$. If no positions existed in the bin, the bin was omitted from the analysis. The allele frequency for the binned data $r_{j'}$ was equal to the mean of the $r_j$ for $j \in \mathcal{J}_{j'}$. The binned data is shown in Fig. 3.

We found that at 250 bp bin size, there was still a significant amount of salt-and-pepper noise: *i.e.* extremely high allele frequency at single isolated position, surrounded by background level activity. Presumably the source of this noise are jackpot events early in proliferation.

To eliminate the jackpot features we use a standard approach from image processing [7]: We generated median filtered allele frequency $\tilde{r}_{j'}$ by taking a median of $r_{j'}$ using the neighborhood $[j'-1, j', j'+1]$. If $r_{j'}$ was four standard deviations above the median-filter value $\tilde{r}_{j'}$, we replace $r_{j'}$ with the median-filtered value $\tilde{r}_{j'}$. The binned and filtered data is shown in Fig. 3.

Note that this binned data $D' = \{(x_{j'}, r_{j'})\}$ is used only in peak detection and the raw (unbinned and unfilter) data $D = \{(x_i, r_i)\}$ is used for model parameter refinement.

##### 2. *Alternative approach*

Some readers may object to the filtering approach we have described. An alternative strategy is to leave the SNPs in the data and use a statistical test to identify them later. In practice, this approach is much slower since it involves optimizing parameters at SNP positions before eventually throwing out these features later. After trying both approaches, we advocate the filtering approach since it produces the same results with much less effort.

##### 3. *Implementing the locus-dependent variance in the test statistic*

Since the dataset is dominated by peak-free regions, we can write:

$$\sigma_i^2 \approx \sigma_0^2 \frac{\mu_i}{\mu_0}, \tag{9}$$

where $\mu_0$ are $\sigma_0^2$ and mean and variance over the entire dataset and $\mu_i$ is the locus-dependent mean (Eq. 1). In this case we know that the tails of the distribution away from the peaks peaks look exponential and not Gaussian. (See Fig. 3B.) If we force the likelihood to be Gaussian, this will inflate the p values. It is therefore convenient to implement the variance model in the following

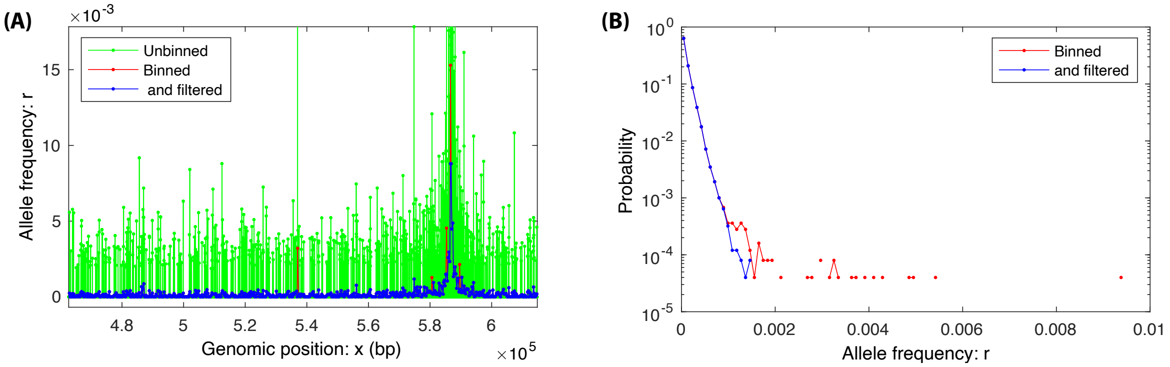


FIG. 3. **Data processing for peak detection for GacA. Panel A:** The raw data (green) was first binned to 250 bp bins (red). Salt-and-pepper noise, isolated high values, were then removed to generated filtered data (blue). Peak detection was performed on the binned and filtered data (blue). Final parameter fits were performed on unbinned and unfiltered data (green). **Panel B:** A histogram of allele frequencies $r_i$. Very few allele frequencies were removed by filtering.

way:

$$\sigma_i^2 \to \sigma_0^2 \max(1, r_i/\mu_0). \tag{10}$$

If the $r_i \gg \mu 0$, the term in the exponent will now be linear:

$$\frac{1}{2\sigma_i^2}(r_i - \mu_0)^2 \approx \frac{1}{2\sigma_0^2} r_i \mu_0, \tag{11}$$

rather than quadratic:

$$\frac{1}{2\sigma_0^2}(r_i - \mu_0)^2 \approx \frac{1}{2\sigma_0^2} r_i^2. \tag{12}$$

This linear dependence matches the observed distribution which decays exponentially in $r$. (See Fig. 3B.) We will use this approach for estimating the p value using the binned data for peak detection.

Another approach will be implemented for parameter approximation (Sec. IV.)

###### 4. *Parameter maximum likelihood estimation for peak detection*

The first step in the null hypothesis test is to perform a maximum likelihood estimate (MLE) of the parameter values. Since the peaks constitute a negligible fraction of the sequence, we will estimate the background mean $\mu_0$ and variance $\sigma_0^2$ using the MLE analysis in the null hypothesis and leave these fixed in all nested models. In what follows, parameters will refer only to the parameters describing the peak profiles. Each peak $J$ will be described by $\theta_J$.

To estimate the parameters from the peak profile, we must first write the minus-log-likelihood for the normal model at the $N$ positions:

$$-\log q(D|\theta) = \frac{N}{2}\log 2\pi\sigma_i^2 + \sum_{i=1}^{N} \frac{1}{2\sigma_i^2}[r_i - \mu_i]^2, \tag{13}$$

where the position-dependent mean $\mu_i(\theta)$ depends on the model through the peak profile function (Eqs. 1-4) and $\sigma_i^2$ is approximated using Eq. 10. We now need to minimize Eq. 13 with respect to the parameters $\theta$.

One difficulty here is that this statistical problem is singular: As $I \to 0$ the peak position $\ell$ becomes unidentifiable [8]. We therefore must take a brute-force approach to estimating $\ell$. We use the following steps: (i) We considered a reduced sets of positions $\ell \in \mathfrak{X} \equiv \{x_i\}_{i=1\ldots N}$. We exhaustively consider a peak position at each $\ell = x_i$. (ii) The parameters $L$ and $a$ were fairly consistent between peaks since they are determined by gross-level chromatin structure. Therefore in the process of peak *detection*, we will assign all peak the global parameter values $\hat{L} \to 400$ bp and $\hat{a} \to 1.5$. (iii) The final unknown MLE parameter $\hat{I}$ can be estimated easily since $\delta\mu_i \propto I$ and a closed-form expression can be derived for it. Since $C'$ has only local support, the MLE estimates can be computed rapidly.

###### 5. *Null hypothesis testing*

The test statistic $\lambda$ in the likelihood ratio test is

$$\lambda(D) \equiv \log \frac{q_1(D|\hat{\theta})}{q_0(D)}. \tag{14}$$

We use the canonical Neyman-Pearson approach to hypothesis testing [1]. We chose a confidence level of $\gamma = 95\%$ (*i.e.* a significance level of 5%). The peak exists, *i.e.* we will reject the null hypothesis, if

$$F_\Lambda(\lambda) > \gamma, \tag{15}$$

where $F_\Lambda$ is the Cumulative Distribution Function (CDF) of the test statistic $\lambda$ under the null hypothesis. (Note that $\lambda$ is capitalized because it is being interpreted as a random variable.)

##### 6. Bootstrap estimate of $F_\Lambda$

Under the normal course of events, if the model were regular in the large sample size limit, we could use the Wilk's theorem to relate the distribution of $\Lambda$ under the null hypothesis to a chi-squares distribution [9]. However, the model is singular [8] and we must therefore estimate the distribution of the test statistic explicitly.

To compute the distribution of the test statistic, we use a stochastic simulation of the null hypothesis and then compute the empirical distribution of the test statistic. Initially we attempted to use a Gaussian random variable to simulate the null hypothesis data, however the estimated p values were too small. In retrospect, it is pretty clear from Fig. 3B that the $r$-distribution tails decay exponentially and therefore large $r_i$'s are much more frequent than predicted by a Gaussian distribution.

In this situation, one can uses a bootstrap method to estimate the test statistic [10]. There are two tractable choices: (i) a the canonical bootstrap approach samples from the empirical distribution consisting of the finite set of observed background allele frequencies shown in Fig. 3B. (ii) A parametric bootstrap method fits the observed distribution to an empirical model and then uses the model to generate simulated data. We used the parametric bootstrap since it had the ability to sample even-more-extreme allele frequencies than were observed. We fit the the distribution of Allele frequencies $r_i$ for the background for the GacA data to the empirical model for random variable $R$:

$$\begin{aligned} p_R(r) &= P_0 \cdot \delta(r) + \ldots \\ &+ \Theta(r)\Theta(r_0 - r) \cdot (1 - P_0)\tfrac{b_1 b_2}{b_+ b_2} e^{-b_1(r_0 - r)} + \ldots \\ &+ \Theta(r - r_0) \cdot (1 - P_0)\tfrac{b_1 b_2}{b_+ b_2} e^{-b_2(r - r_0)}, \end{aligned} \tag{16}$$

where $\delta$ and $\Theta$ are the Dirac delta and Heaviside function respectively. See Fig. 4. The empirical model parameters were fit using an MLE approach:

$$r_0 = 7.2906 \times 10^{-5}, \tag{17}$$

$$P_0 = 5.386 \times 10^{-1}, \tag{18}$$

$$b_1 = 1.2972 \times 10^{5}, \tag{19}$$

$$b_2 = 8.4846 \times 10^{3}. \tag{20}$$

The fit of the empirical model to the background allele frequency is excellent and is shown in Fig. 4.

##### 7. Estimating the distribution of the test statistic

Using the parametric-bootstrap model, we simulated the null hypothesis data $D' = \{(x_{j'}, R_{j'})\}$ where $R_{j'} \sim p_R$. For each simulated dataset, we then computed the test statistic:

$$\Lambda \equiv \lambda(D'), \tag{21}$$

which we interpret as a random variable. We generated $10^5$ samples of $\Lambda$. We then use the empirical distribution of $\Lambda$ to estimate the p values in the usual way (e.g. [10]).

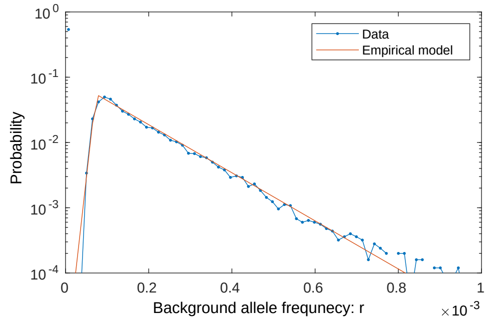


FIG. 4. **Empirical model for parametric bootstrap.** The measured background allele frequency is shown in blue and the empirical model fit by MLE analysis is shown in red.

##### 8. Computation of the p value

We have included a p value for each detected peak as a proxy for statistical support. The p value for test statistic $\lambda$ is:

$$p(\lambda) = 1 - F_\Lambda(\lambda). \tag{22}$$

Since some of peaks are extremely large, the observed test statistic is much larger than any observed in our simulations. To estimate the p values in this context, we fitted the empirical distribution $F_\Lambda$ to a Gumbel distribution since the minimization of the minus-log-likelihood over $\ell$ can be reinterpreted as an extreme value problem for a random variable in the exponential family [11]. The Gumbel distribution is

$$F_\Lambda(\lambda) = e^{-e^{-\frac{\lambda - \mu_\Lambda}{\sigma_\Lambda}}}, \tag{23}$$

where the position and scale parameters are

$$\mu_\Lambda = 0.7217, \tag{24}$$

$$\sigma_\Lambda = 0.2823, \tag{25}$$

respectively, which we estimated using an MLE approach. For very small $p$ we can make the following approximation:

$$\log p \approx -\tfrac{\lambda - \mu_\lambda}{\sigma_\lambda}, \tag{26}$$

by Taylor expanding the outer-most exponential around zero in Eq. 23.

##### 9. Statistical tests for subsequent nested models

After a peak is detected by rejecting the null hypothesis, we replace the null hypothesis with the alternative hypothesis and then define a new alternative hypothesis with another putative peak. We then repeat the null hypothesis test. This procedure was repeated until no more statistically significant peaks could be detected.

#### IV. PARAMETER INFERENCE AND FIT REFINEMENT

Once the peaks were detected, we refined all four profile parameters, $\theta = (I, \ell, a, L)$, for each peak by direct numerical maximum likelihood estimation for all parameters, now all defined on $\mathbb{R}$. Note that this optimization is performed *after* peak detection. This refinement is performed on the full resolution data.

For parameter inference we will use a different approach for the scaling of the variance:

$$\sigma_i^2 = \sigma_0^2 \frac{\mu_i}{\mu_0}, \tag{27}$$

since the approximation in Eq. 10 fails for the higher resolution data. For parameter optimization, the tails of the distribution are of little importance.

To estimate the uncertainty in the parameters, we used the Fisher information in the usual way (e.g. [1]). The numerical minimization resulted in a Jacobian:

$$J_{\alpha i} \equiv [\partial_{\theta_\alpha} \delta\mu_i / \sigma_i, ](\hat{\theta}). \tag{28}$$

The Fisher information is then:

$$I = [J J^T], \tag{29}$$

and therefore the predicted covariance in error is

$$\overline{\delta\theta_\alpha \delta\theta_\beta} = [I^{-1}]_{\alpha\beta}. \tag{30}$$


[1] D. R. Cox and D. V. Hinkley, *Theoretical Statistics* (Chapman & Hall, 1974).
[2] K. Rippe, P. H. von Hippel, and J. Langowski, Trends in Biochemical Sciences **20**, 500 (1995).
[3] J. Dekker, K. Rippe, M. Dekker, and N. Kleckner, Science **295**, 1306 (2002).
[4] E. Lieberman-Aiden, N. L. van Berkum, L. Williams, M. Imakaev, T. Ragoczy, A. Telling, I. Amit, B. R. Lajoie, P. J. Sabo, M. O. Dorschner, R. Sandstrom, B. Bernstein, M. A. Bender, M. Groudine, A. Gnirke, J. Stamatoyannopoulos, L. A. Mirny, E. S. Lander, and J. Dekker, Science **326**, 289 (2009).
[5] S. E. Luria and M. Delbrück, Genetics **28**, 491 (1943).
[6] Q. Zheng, Mathematical Biosciences **162**, 1 (2010).
[7] R. C. Gonzalez and R. E. Woods, *Digital image processing* (Prentice Hall, Upper Saddle River, N.J., 2008).
[8] S. Watanabe, Journal of Machine Learning Research. **14**, 867 (2013).
[9] S. S. Wilks, The Annals of Mathematical Statistics. **9**, 60 (1938).
[10] B. Efron and R. Tibshirani, *An Introduction to the Bootstrap.* (Chapman & Hall/CRC, Boca Raton, FL, 1993).
[11] L. Haan and A. Ferreira, *Extreme value theory: an introduction.* (Springer., 2007).
